# Genetic factors explaining anthocyanin pigmentation differences

**DOI:** 10.1101/2023.06.05.543820

**Authors:** Maria F. Marin Recinos, Boas Pucker

**Affiliations:** Plant Biotechnology and Bioinformatics, Institute of Plant Biology and BRICS, TU Braunschweig, Braunschweig, Germany

**Keywords:** Flavonoid biosynthesis, Anthocyanins, Gene expression, Transcription factor, MYB, Systematic comparison, Transcriptome

## Abstract

**Background:** Anthocyanins represent one of the most abundant coloration factors found in plants. Biological functions of anthocyanins range from reproduction to protection against biotic and abiotic stressors. Owing to a clearly visible phenotype of mutants, the anthocyanin biosynthesis and its sophisticated regulation have been studied in numerous plant species. Genes encoding the anthocyanin biosynthesis enzymes are regulated by a transcription factor complex comprising MYB, bHLH and WD40 proteins.

**Results:** A systematic comparison of anthocyanin-pigmented vs. non-pigmented varieties across flowering plant species was performed. Literature was screened for cases in which genetic factors causing anthocyanin loss were reported. Additionally, transcriptomic data sets from previous studies were reanalyzed to determine the genes most likely to be responsible for color variation based on their expression pattern. The contribution of different structural and regulatory genes to the pigmentation differences was quantified. Gene expression differences concerning transcription factors are by far the most frequent explanation for pigmentation differences observed between two varieties of the same species. Among the transcription factors in the analyzed cases, MYB genes are substantially more likely to explain pigmentation differences than bHLH or WD40 genes.

**Conclusions:** These findings support previous assumptions about the plasticity of transcriptional regulation and its importance for the evolution of novel coloration phenotypes. Our findings underline the particular significance of MYBs and their apparent dominant role in the specificity of the MBW complex.

## INTRODUCTION

Angiosperms are characterized by an enormous diversity of flower hues and shapes[1,2]. Some plant species maintain their brilliant colors throughout the year, while others show a constant transformation as the seasons change. The substances responsible for these colors are pigments which include flavonoids, betalains, and carotenoids[3]. These pigment classes differ in their biochemical properties resulting in distinct color ranges. Flavonoids can be classified into multiple subgroups with anthocyanins forming the most colorful subgroup. Anthocyanins can provide orange, red, purple, blue, or almost black coloration[4]. Carotenoids lead to yellow, orange, or red coloration[5]. Betalains can be classified into yellow betaxanthins and red betacyanins[3].

Anthocyanins and other flavonoids are a group of specialized plant metabolites responsible for numerous functions beyond coloration. Additional physiological functions are protection against herbivores[6,7] and reduction of the impact caused by salinity[8], drought[9], and UV-radiation[10,11]. Associated with their color are ecological functions such as the attraction of pollinators and seed dispersers which facilitates reproduction[6,12]. Not only anthocyanins, but also flavonols are contributing to the attraction of pollinators by forming guiding signals on flowers which are invisible to the human eye[12,13]. Pigmented anthocyanins are known to be responsible for the coloration of flowers ranging from red to orange to purple and to blue colors. The diversity in colors depends on the chemical structure of the anthocyanin compound which includes the number of hydroxyl groups attached to the benzene ring, and the level of glycosylation[14,15] and acylation[16]. Several reports suggest that the interaction with copigments like flavonols and flavones is an important factor for the stabilization of anthocyanins in plants[17,18]. Environmental factors can also influence the color stability of anthocyanin pigments, for example, a plant exposed to an acidic soil can produce anthocyanins with an intense red or orange color[19], plants exposed to high temperatures can show degradation of anthocyanins while low temperatures can increase color intensity[13].

Besides anthocyanins there are three major classes of flavonoids produced by a wide range of plant species: flavones, flavonols, and proanthocyanidins. Each of these subgroups of flavonoids have individual biological functions and can influence the coloration of different plant structures[20]. The characteristic cream white or pale yellow color which determined the presence of flavones and flavonols can be observed in leaves or petals of *Taraxacum officinale* “dandelion”[21], *Chrysanthemum grandiflorum* cv. Jinba[22], and *Chrysanthemum morifolium*[23]. Proanthocyanidins also called condensed tannins are colorless compounds that turn brown upon oxidation[24]. They have been studied in seeds of species such as *Arabidopsis thaliana*[25], *Brassica napus*[26], and *Ipomoea purpurea*[27].

The general pathway of the flavonoid biosynthesis (Figure 1A) is well understood and the central aglycon biosynthesis is largely conserved among land plants[3,28]. It starts with the condensation of 4-coumaroyl-CoA and malonyl-CoA to synthesize naringenin chalcones which are later isomerized by the enzyme chalcone isomerase (CHI) to form naringenin, a colorless flavanone. In the next step, the pathway diverges: flavanone can either be hydroxylated by the flavanone-3-hydroxylase (F3H) to form dihydroflavonols or it can be oxidated through the activity of flavone synthase (FNS) and form flavones. After the hydroxylation of naringenin to dihydrokaempferol, the formation of dihydroquercetin and dihydromyricetin can take place through the catalysis of flavonoid 3’-hydroxylase (F3’H) and flavonoid 3’,5’-hydroxylase (F3’5’H), respectively. Subsequently, two enzymes can accept these intermediates and produce either flavonols through the oxidation with flavonol synthase (FLS) or leucoanthocyanidins by the reduction with dihydroflavonol 4-reductase (DFR). In case the latter occurs, another enzyme called anthocyanidin synthase/leucoanthocyanidin dioxygenase (ANS/LDOX) catalyzes the synthesis of anthocyanidins. This step also requires an anthocyanin-related gluthationine S-transferase (arGST) that was named AN9/TT19 due to the corresponding mutants[29,30], but the enzymatic function was only revealed recently [31]. These anthocyanidins can be further modified through different steps including (Figure 1B) (1) glycosylation in the presence of UDP-glucose flavonoid 3-O-glucosyl transferase (UFGT), (2) methylation through the activity of O-methyltransferase (OMT), and (3) acylation by the anthocyanin acyltransferase (ACT). Moreover, leucoanthocyanidins and anthocyanidins can also be reduced by the enzymatic activity of leucoanthocyanidin reductase (LAR) and anthocyanidin reductase (ANR), respectively, to synthesize catechins and epicatechins leading to the production of proanthocyanidins.

**Figure 1.**
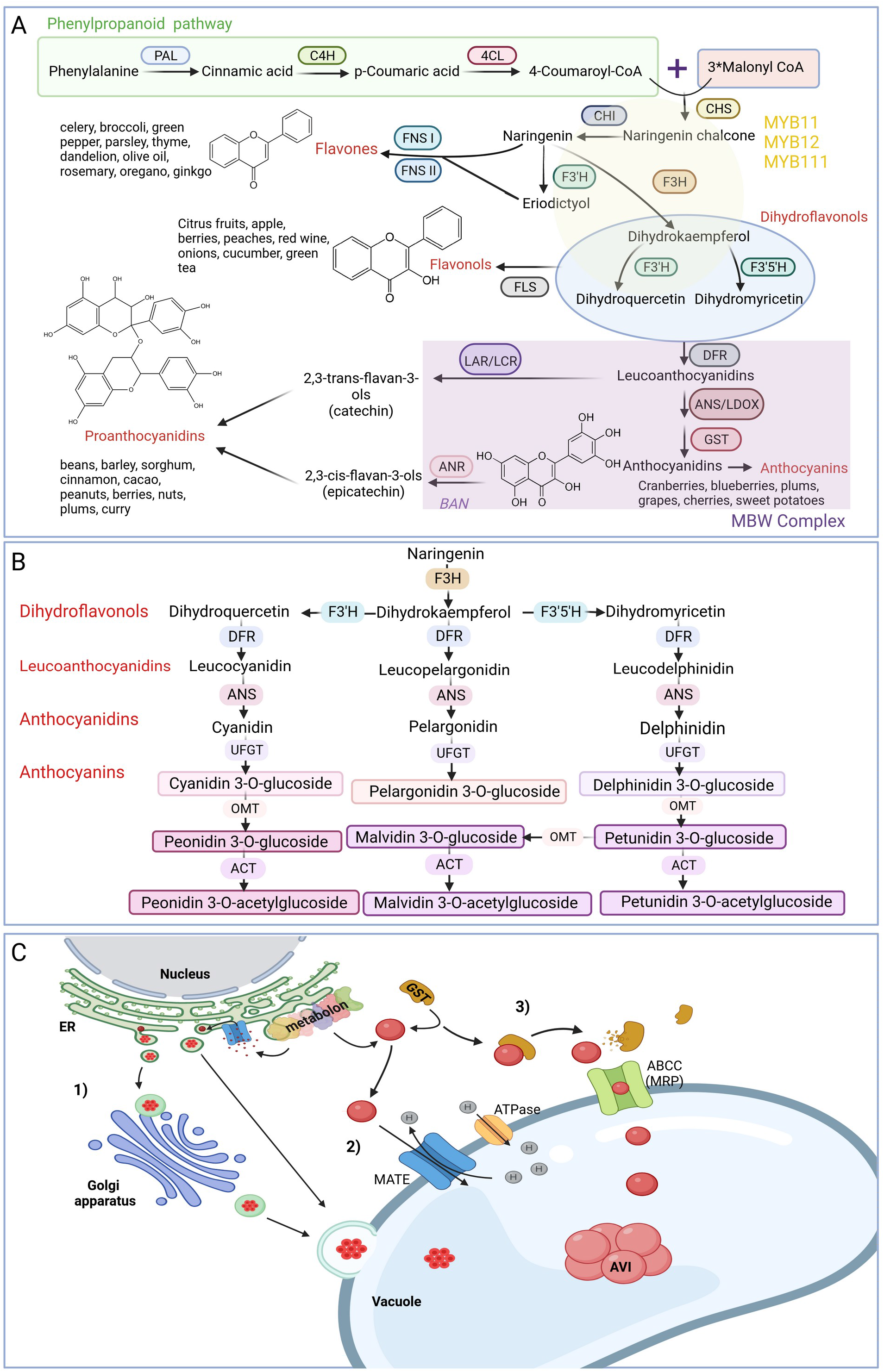
(A) Schematic representation of the general flavonoid biosynthesis pathway. (B) Simplified flowchart describing the biosynthesis pathway of anthocyanins. Enzyme names are abbreviated as follows: PAL – phenylalanine ammonia-lyase, C4H – cinnamic acid 4-hydroxylase, 4CL – 4-coumarate-CoA ligase, CHS – chalcone synthase, CHI – chalcone isomerase, F3H – flavanone 3-hydroxylase, F3’H – flavonoid 3’-hydroxylase, F3’5’H – flavonoid 3’,5’-hydroxylase, DFR – dihydroflavonol 4-reductase, ANS – anthocyanidin synthase, UFGT UDP-glucose:flavonoid 3-O-glucosyltransferase, FLS – flavonol synthase, ANR – anthocyanidin reductase, ANS-Anthocyanidin syntase, LDOX – leucoanthocyanidin dioxygenase, FNS – flavone synthase. (C) Scheme of the different intracellular flavonoid transport mechanisms. 1. vesicle trafficking from the Endoplasmic Reticulum (ER) 2. membrane transporter throughout the Multidrug and Toxin Extrusion Transporter (MATE), and 3. Transport might be mediated by the Glutathione S-transferase (GST) and the ATP-binding cassette (ABCC) transporter. GST is presented twice due to the recently reported enzymatic function of arGSTs by Eichenberger *et al.*[31]. It is currrently not clear if GST functions only as enzyme or if they also play a role in anthocyanin transport.

Like many other specialized metabolites, flavonoids are synthesized in the cytoplasm. It is assumed that some enzymes responsible in catalyzing specific reactions in the flavonoid biosynthesis are attached to the endoplasmic reticulum (ER) and form a metabolon[32](Figure 1C). Flavonoids produced at the ER are transported into the vacuole for storage[33–35] which results in observable pigmentation.

The transport of these metabolites to the vacuole is a process that is not fully understood, but different models have been proposed that could explain observations of a range of experiments. Two widely accepted and well-known models are: (1) vesicle trafficking from the ER to the vacuole and (2) GST-mediated transport to the tonoplast, where membrane-based transporters are active[34]. Both models have in common that a mechanism is required to transport flavonoids across a membrane and these models are not mutually exclusive. It is feasible that these transport routes are active to certain degrees under different conditions, in developmental stages, or in different plant parts. The vesicular transport model proposes the accumulation of flavonoids inside the ER lumen and formation of small flavonoid rich compartments surrounded by a membrane that move to the central vacuole[32,35]. The existence of these vesicles has been reported mainly in *Zea mays*[36], *A. thaliana*[37], *Vitis vinifera*[38], and *Oryza sativa*[39]. Microscopic evidence shows that these vesicular bodies are attached to the surface of the ER[38,40] from where they are released into the cytoplasm and mobilized directly into the vacuole either by fusing with carrier proteins, or mobilized indirectly by following the trans-Golgi Network (TGN) transport pathway[41]. The GST-mediated transport model proposes that flavonoids are delivered to the tonoplast by ligandins[29,42]. These ‘ligandins’ would be glutathione S-transferase (GST) binding and carrier proteins[43]. Evidence for the role of GST in flavonoid transport has been reported in multiple species, such as *Zea mays*[42], *A. thaliana*[29,30]*, Petunia hybrida*[43]*, Vitis venifera*[38], and *Prunus persica*[44]. Reconsideration of the GST-mediated transporter is needed in the light of a recent study[31] that revealed an enzymatic function in the anthocyanin biosynthesis for anthocyanin-related GSTs (arGSTs). The tonoplast-based transport mechanism involves different transmembrane channels which enable translocation of flavonoids into the vacuole[32]. These routes were reported to involve the Multidrug Resistance-associated Protein (MRP), belonging to the family of proteins ATP-binding cassette (ABC) actively transporting anthocyanins[45]. Using an electrochemical H^+^ gradient to transport substances across membranes, Multidrug and Toxic compound Extrusion (MATE) is considered to regulate the vacuolar sequestration of proanthocyanidin precursors in the seed coat cells[45–47].

The transcriptional activity of genes encoding enzymes of the flavonoid biosynthesis pathway is controlled by numerous transcription factors or even protein complexes comprising multiple transcription factors (Figure 1A). Genes of the flavonol and flavone biosynthesis are largely regulated by MYB11, MYB12, and MYB111[48–50]. The mechanism that regulates the expression of all structural genes in the anthocyanin biosynthesis pathway is commonly known as the MBW complex[51]. The name of the MBW complex is based on the three involved transcription factors: R2R3-MYBs, basic helix-loop-helix (bHLH) proteins, and WD40 proteins. One member of each of the three protein families is required for the complex formation. Different members of the MYB and bHLH family can participate resulting in combinatoric diversity[51]. After the discovery of TT8 in *A. thaliana*[52], Baudry *et al*.[53] demonstrated the activity of the MBW complex in regulating the expression of the proanthocyanidins (PA) biosynthesis gene *BANYULS* (*BAN*). The ternary complex responsible for *BAN* regulation in *A. thaliana* is composed of TT2 (MYB123), TT8 (bHLH42), and TTG1 (WD40 family). Years later, it was demonstrated that the anthocyanin biosynthesis is also controlled by MBW complexes[54]. These complexes harbor PAP1, PAP2, MYB113, or MYB114 as the MYB component and GL3 or EGL3 as bHLH component as well as the WD40 protein TTG1[54].

It is assumed that the function of regulating anthocyanin biosynthesis by the MBW complex is evolutionary conserved across angiosperms[55]. Elements of the MBW complex are also involved in the control of other biological processes in plants[56]. Other functions add different degrees of evolutionary constraints to the components of the MBW complex. MYB partners are substituted and considered as the specificity determining factor in the MBW complex. For example, PAP1/MYB75 and PAP2/MYB90 activate the anthocyanin biosynthesis[57], while TT2/MYB123 would activate the proanthocyanidin biosynthesis[53]. The DNA binding and protein-protein interaction capacity of MYBs and bHLHs is determined by highly conserved regions[58]. It has been postulated that TTG1 serves as a scaffolding protein that maintains the interaction of MYB and bHLH[59]. This protein-protein interaction involves five WD repeats that account for over 60% of the protein length. Previous reports suggest that the MYB component is most often associated with changes in flower pigmentation indicating low constraints on this component of the MBW complex due to higher functional specialization[60–62].

Closely related plant species can differ in their anthocyanin repertoire and pigmentation pattern[63,64]. These differences can even appear between plants of the same species[1,65–67]. All structural genes in the anthocyanin biosynthesis pathway must be functional and active to achieve anthocyanin pigmentation. Mutations in any regulator or enzyme encoding gene of the flavonoid biosynthesis can affect the coloration. Pigmentation differences have been studied in many plant species including *A. thaliana*[68], *Vitis vinifera*[69], *Hordeum vulgare*[70], *Nicotiana tabacum*[71], and *Nicotiana alata*[72]. Substrate competition between different branches of the flavonoid biosynthesis can also have an impact on the anthocyanin accumulation[73]. For example, an increased flavonol production can lead to a pigmentation loss due to insufficient substrate for the anthocyanin biosynthesis[74]. These natural differences in pigmentation provide an excellent system to study evolutionary processes that lead to the inactivation of a pathway. In theory, a biosynthesis pathway could be interrupted at any of the successive steps[75], but previous research suggests that some genes are more often responsible for pigmentation loss than others[62,76]

This raises the question whether there are “hot spots” in the pathway that are frequently the cause for a pigmentation loss; not due to higher mutation rate, but higher likelihood of mutation fixation. Specifically, we ask three questions: (1) Are anthocyanin pigmentation differences within species caused mainly by transcription factors or structural genes? (2) Is *DFR* at the start of the anthocyanin biosynthesis branch more likely to be causal for anthocyanin variations than downstream genes like *ANS*, *arGST*, or *UFGT*? (3) Are MYB more often responsible for pigmentation differences than other transcription factor families? To address these questions, we performed a systematic comparison of anthocyanin-pigmented vs. non-pigmented varieties across flowering plant species. The literature was screened for reports of causal genes explaining pigmentation differences between varieties of a plant species or between closely related (sub)species. Studies without conclusive results were checked for data availability and subjected to reanalysis when possible. The contribution of different genes to the pigmentation differences was quantified and supported a crucial role of transcription factors and especially MYBs.

## RESULTS

### Genetic hot spots responsible for anthocyanin pigmentation differences

Based on a total of 230 analyzed studies (see Additional file1: Table S1) including reanalyzed RNA-seq experiments, we determined the genes most likely to be responsible for color variation between accessions in each of these species. In our schematic representation (Figure 2), we defined pigmentation to be the wild type state, while absence of pigmentation was defined to be the result of a mutation. We identified 13 research articles that report upregulated structural genes in pathways competing for substrate with the anthocyanin biosynthesis as the cause of color difference between unpigmented and pigmented accessions. Additionally, 58 events of non-activated, down-regulated, non-functional, or lost structural anthocyanin biosynthesis genes were reported in the literature. Moreover, in 147 different cases a transcription factor was proposed to be responsible for differences in pigmentation. Many of these reports named a specific transcription factor. In total, 82 MYBs (activators and repressors), 10 bHLHs, two TTG1 homologs, one bZIP, one WRKY were presented as the causal gene for pigmentation differences. The remaining 49 cases are likely due to the action of multiple transcription factors or caused by TFs that activate the components of the MBW complex. Ten reports presented genes that encode proposed intracellular transporters of anthocyanins as best candidates such as MATE and possibly GST. It was not possible to determine the causal gene in 17 of the analyzed studies.

**Figure 2.**
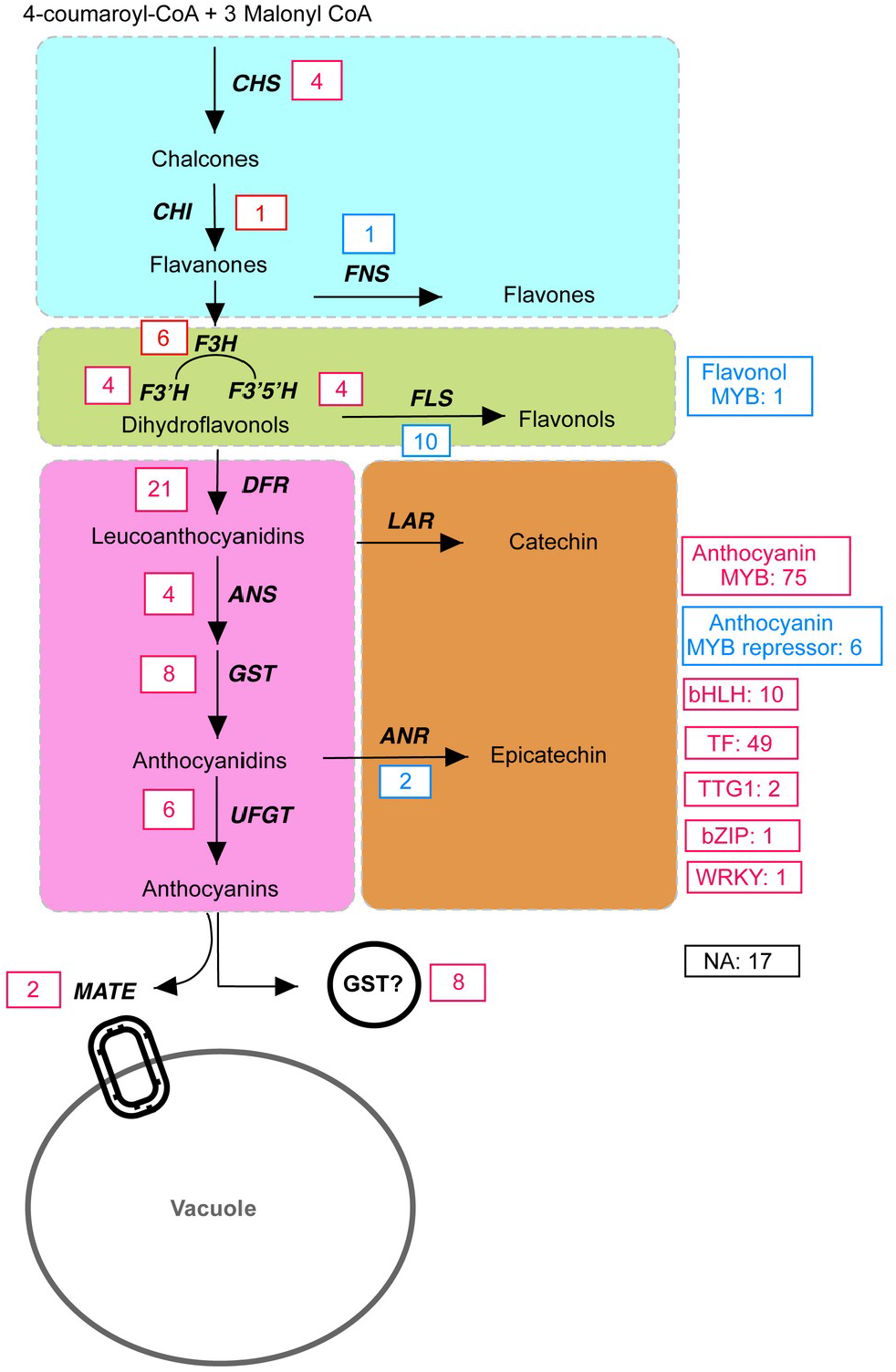
Schematic representation of the flavonoid biosynthesis pathway with emphasis on the number of cases in which a particular gene was responsible for color difference according to a systematic literature analysis and re-analyses of RNA-seq data sets. The anthocyanin-pigmented accession is set as reference when determining up- and down-regulation. Blue boxes and red boxes indicate the number of upregulated and downregulated/non-functional/lost genes, respectively. Downregulated genes are placed in one group with genes that lost their function due to mutations in the coding sequence or completely lost genes, because the ultimate function of the gene is lost in any of these cases. Flavonoids were divided into four groups that are indicted by color shading: flavones, flavonols, anthocyanins, and proanthocyanidins. Black bold letters represent the different genes encoding the enzymes and transporters involved in the pathway. *CHS* – chalcone synthase, *CHI* – chalcone isomerase, *F3H* – flavanone 3-hydroxylase, *F3’H* – flavonoid 3’-hydroxylase, *F3’5’H* – flavonoid 3’,5’-hydroxylase, *DFR* – dihydroflavonol 4-reductase, *ANS* – anthocyanidin synthase, *UFGT* – UDP-glucose:flavonoid 3-O-glucosyltransferase, *FLS* – flavonol synthase, *LAR* – leucoanthocyanidin reductase, *ANR* – anthocyanidin reductase, FNS – flavone synthase, GST – Glutathione S-tranferase, MATE – Multidrug And Toxin Extrusion, TF – unclassified transcription factor. GST is presented twice due to the recently reported enzymatic function of arGSTs by Eichenberger *et al.* [31]. It is currrently not clear if GST functions only as enzyme or if they also play a role in anthocyanin transport.

### Taxonomic distribution of analyzed cases across plant families

The 230 reviewed studies (see Additional file 1: Table S1), cover 53 plant families distributed over 30 orders (Figure 3). Notably, the order Rosales, particularly the Rosaceae family, accounted for the highest number of studies (39) showcasing variations in pigmentation. Second were the Brassicaceae family with 24 reported cases, and the Orchidaceae, Fabaceae, and Solanaceae families each contributing 13 cases. Furthermore, the Ericaceae family appearing in 9 studies, and the Asteraceae and Liliaceae families, each appearing in 8 studies, were also noteworthy. While the Theaceae, Poaceae, and Paeoniaceae families were featured each in 7 cases. Other families, including Lamiaceae, Asparagaceae, Malvaceae and Caryophyllaceae, were identified in varying frequencies across the reviewed studies. This rich diversity in distribution of families and orders is crucial to reveal universal mechanisms explaining pigmentation differences within plant species. Additionally, it highlights the ecological and evolutionary significance of this morphological phenomenon across angiosperms. However, it is noteworthy to clarify that among the different studies the terms “varieties”, “lines”, and “cultivars” were often used interchangeably when comparing intraspecific plants. In the literature, it was not consistently clarified whether these distinctions arose from horticultural/artificial interventions, or if those differences could be attributed to natural causes. This nuance would be helpful for the interpretation of the results to distinguish between naturally fixed variants and those selected by breeders.

**Figure 3.**
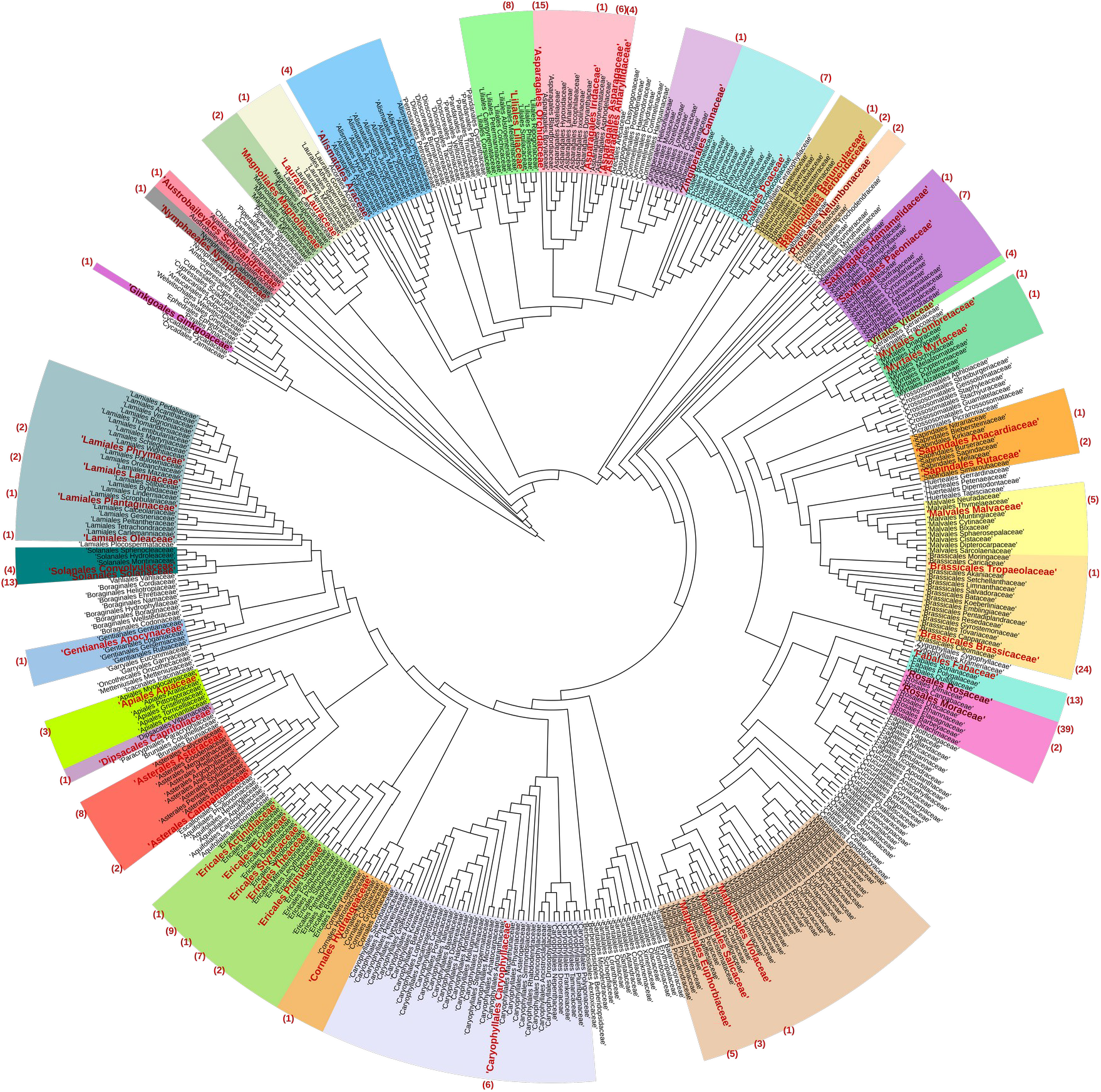
Phylogenetic tree displaying 428 angiosperm families. Each color range groups the families of an order. Families highlighted in bold red are those encompassed in the literature screening for anthocyanin pigmentation differences. The number of pigmentation difference cases is given in parentheses for each family. Modified from Li *et al.*[77].

### Genetic factor of anthocyanin loss across plant families

The genes that have been identified as influential factors in driving variation were graphically represented along with the families in which they have been reported (Figure 4). This visualization aims to uncover potential associations between specific genes and their prevalence across different plant lineages.

A noteworthy observation is the prominence of anthocyanin biosynthesis activating MYB transcription factor genes (classified as AnthoMYBact), which have been reported in 75 cases across all plant families. Families with a particular high prevalence of AnthoMYBact are Rosaceae, Brassicaceae, Orchidaceae, Liliaceae, Solanaceae, and Asteraceae. On the contrary, some genes such as anthocyanin biosynthesis repressing MYBs (AnthoMYBrep), transcription factor bZIP, enzyme CHI, and FlavonolMYB appear in fewer instances, indicating a rare involvement in color variation. By juxtaposing genes causal for anthocyanin loss with the respective families where they have been observed, we aim to discern any family-specific patterns.

**Figure 4.**
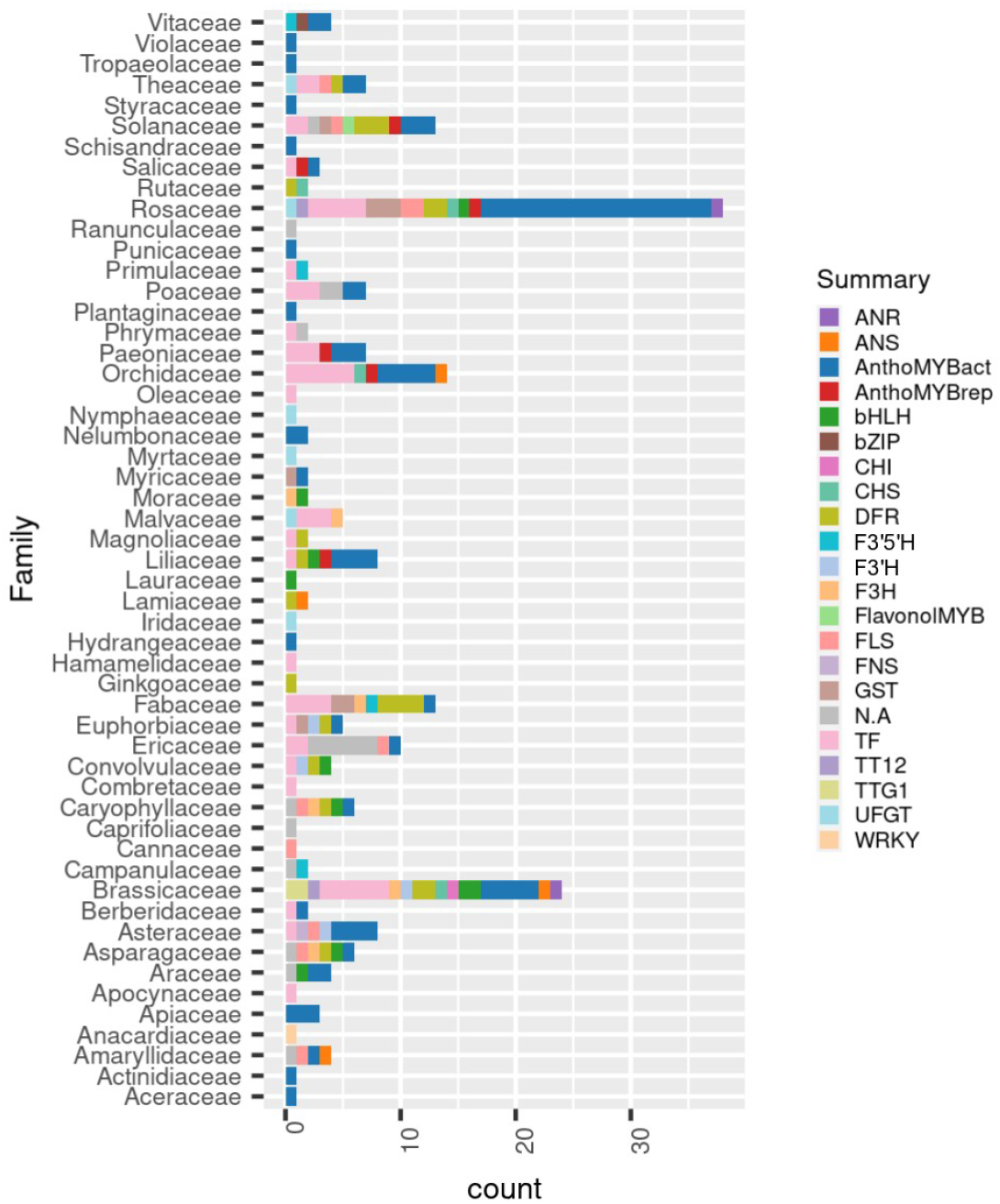
Number of cases each gene is implicated in anthocyanin differences reported in the literature (Additional file 1: Table S1) resolved by family. ANR – Anthocyanidin reductase, ANS – Anthocyanidin synthase, AnthoMYBact – Anthocyanin MYB activator, AnthoMYBrep – Anthocyanin MYB repressor, bHLH – Basic Helix-Loop-Helix, bZIP – Basic leucine zipper, CHS – Chalcone synthase, CHI – Chalcone isomerase, DFR – dihydroflavonol 4-reductase, F3’5’H – flavonoid 3’,5’-hydroxylase, F3’H – flavonoid 3’-hydroxylase, F3H – flavanone 3-hydroxylase, FLS – flavonol synthase, FNS – flavone synthase, GST (arGST) – anthocyanin-related glutathione S-transferase, TF – Transcription factor, TT12 – Transparent Testa 12, TTG1-Transparent Testa Glabra 1, UFGT – UDP-glucose:flavonoid 3-O-glucosyltransferase, WRKY-WRKY DNA-binding domain, N.A – Unknown factor.

### Overrepresentation of genetic factors causing anthocyanin loss

The hypothesis that *DFR* might be more often responsible for an anthocyanin loss than *ANS* or other players in the anthocyanin biosynthesis was tested. Among the 230 studies analyzed (see Additional file 1: Table S1), *DFR* exhibited the highest frequency, being reported in 21 cases, while *ANS* was identified in four cases as the gene responsible for color variation. The number of cases reporting *DFR* as the primary factor for pigmentation differences is significantly higher than the number of cases reporting *ANS* (χ² test, p-value = 0.00067).

To further examine whether this dominance of *DFR* extends to subsequent anthocyanin genes, we compared *DFR* with *arGST* which also revealed a significant difference (χ² test, p-value = 0.0158). Similarly, when examining the relationship between *DFR* and *UFGT*, a notable difference was again detected (χ² test, p-value = 0.0039). This suggests that *DFR* is the most important target of evolutionary events blocking the anthocyanin pathway.

To investigate if the color variation could be attributed to the substrate competition between FLS and DFR, we examined the prevalence of cases revealing *FLS* and *DFR* as the primary genes influencing color variation in plant tissue. A hyperactivation of *FLS* was identified in 10 cases as the factor responsible for pigmentation differences which is significantly lower than the number of 21 cases in which a DFR down-regulation/loss was responsible (χ² test, p-value = 0.048). This suggests a more pronounced influence of DFR in the observed color variations, implying a potentially pivotal role in the genetic mechanisms governing anthocyanin production. It further suggests that a downregulation or silencing of *DFR* is more strongly associated with the white coloration of plant tissues compared to an increased activity in expression of an *FLS* gene and subsequent production of flavonols.

Similar to the comparison between the structural genes, the differences between the TFs reported to be the causal factor of anthocyanin pigmentation differences were analyzed. The chi-square analysis revealed that the frequency of anthocyanin biosynthesis activating MYBs appearing as causal gene of color variation is significant higher than the presence of other transcription factor families such as bHLH, WRKY, TTG1, bZIP, and others (χ² test, p-value = 1.61e-10). Even when compared against the large group of unclassified TFs, we observed that the presence of MYBs is significantly higher (χ² test, p-value = 0.019). It is expected that most of the cases with unidentified transcription factors would actually be due to anthocyanin biosynthesis activating MYBs. In summary, these findings collectively highlight the dominant role played by the MYB transcription factor family in influencing the observed pigmentation variations.

## DISCUSSION

Anthocyanins are one of the main factors responsible for color variation in plant tissues, particularly in flowers. The variation in floral coloration can occur as a result of plant adaptation to different biotic and abiotic conditions, but interactions with pollinators might be the most important function of anthocyanins in flowers e.g. to indicate which flowers have already been visited by insects[78]. Flavonoids are also known to protect against UV radiation[79], drought[80], and cold stress[81,82]. Previous studies have reported numerous genes responsible for anthocyanin pigmentation differences within a plant species (Additional file 1: Table S1). We present an aggregated analysis of the most likely candidates genes responsible for color variation. This analysis harnessed numerous public RNA-seq data sets that enable a direct comparison of anthocyanin biosynthesis gene activity between differently pigmented accessions and the taxonomical family they belonged. This comparison evaluates whether specific genes are responsible for pigmentation loss in certain lineages. The complex interplay between specific genes and their role in shaping plant pigmentation has been a subject of investigation in numerous studies[83,73,84–87] and has been experimentally tested in families such as Solanaceae[86], as well as in specific genera like Ipomoea[61,63], Iris[88], Antirrhinum[89], and Petunia[76].

### DFR is the block hotspot in the anthocyanin branch

DFR activates the conversion of dihydroflavonols to leucoanthocyanidins, which is often considered as the first committed step of the anthocyanin biosynthesis. Through a comprehensive literature survey, we revealed that *DFR* is more often harbouring a disruptive mutation that results in a block of the anthocyanin accumulation in colorless varieties than any downstream gene in the anthocyanin biosynthesis. A transition from red/purple to white/cream flower color would require some kind of blockage in the anthocyanin production, which probably occurs upstream of anthocyanidin formation[90]. Leucoanthocyanidins can be catalyzed to form two different products, catechins via LAR and anthocyanidins via ANS. If the anthocyanin biosynthesis is blocked at the ANS step, the product to be formed would be catechins. This redirection in the metabolic flow would form proanthocyanidins instead of anthocyanins and could ultimately result in brownish pigmentation due to polymerized and oxidized proanthocyanidins[91].

A study published by Rausher *et al*.[92] proposed that the evolutionary rate of enzymes depends on their location in a pathway with early genes showing a slower mutation rate, but this has been contradicted recently[93]. According to the Arabidopsis Information Resource (TAIR), the length of the coding sequence (CDS) of *DFR* is 1149 bp (accession: NM_123645) and of *ANS* is 1071 bp (accession: NM_118417). To the best of our knowledge and based on the low length difference between the *DFR* and *ANS* coding sequences, there is no evidence that the occurrence of a mutation in *DFR* is substantially more likely than a mutation in *ANS*. While the mutation rate in both genes is probably equal, the rate of mutation fixation might be very different. According to theories of metabolic regulation, it is evolutionary beneficial to have blocks at the first committed step of a branch in a biosynthesis pathway in order to avoid a waste of energy and resources by pushing substrate into a blocked pathway[94,95]. This could explain why *DFR* and not *ANS*, *arGST*, or *UFGT* appears frequently in analyzed cases of intraspecific anthocyanin pigmentation differences.

### Cross talk and substrate competition: Anthocyanins vs. flavonols

Plants have multiple mechanisms to regulate their metabolism in response to environmental conditions and availability of resources. Substrate competition is among the factors determining the color variation observed in plant tissues[17]. Metabolically, this can occur when two enzymes or transporters compete for the same or very similar substrates[94]. DFR and FLS are both catalyzing reactions that utilize dihydroflavonols, but lead to different products. While DFR generates colorful anthocyanins, FLS produces colorless flavonols. There are three different types of dihydroflavonols namely dihydrokaempferol, dihydroquercetin, and dihydromyricetin that differ in their hydroxylation pattern. Different isoforms of DFR and FLS have preferences for specific hydroxylation patterns which could be a mechanism to avoid direct substrate competition. The relative activities of F3H, F3’H, and F3’5’H determine the intracellular levels of the three dihydroflavonols. Our analyses revealed that variation associated with DFR is more often responsible for a color change than variation associated with FLS. In total, 21 cases revealed that the low expression of *DFR* is responsible for the color contrast between unpigmented tissues and those that show anthocyanin pigmentation. Only ten cases showed an increased FLS activity as the cause of pigmentation loss. A high expression of *FLS* leads to an accumulation of colorless flavonols instead of colorful anthocyanins as reported previously in several plant species[96]. A recent study identified a flavonol biosynthesis regulating MYB as the most frequently affected gene in pigmentation pattern change[93]. Loss of expression or loss of a gene function can be the consequence of many different mutations and thus be more likely to happen than a gain-of-function mutation. It is also feasible that some researchers only investigated the classical anthocyanin biosynthesis genes when looking for a molecular mechanism to explain the anthocyanin pigmentation difference thus leading to an observation bias concerning the responsible genes. However, this is unlikely to explain the strong difference between hotspots like DFR and MYB and other genes. Once the anthocyanin biosynthesis is disrupted, selection against additional mutations in the anthocyanin biosynthesis genes might be weak or completely absent. This could result in the accumulation of secondary mutations. A number of additional mutations in the anthocyanin biosynthesis would increase the chances that at least one of them is picked up by researchers looking at anthocyanin biosynthesis genes. Performing future analyses by inspecting a more comprehensive gene sets could make the identification of causal genes in color difference studies more accurate.

### Transcription factor variations appear frequently as block to anthocyanin accumulation

It is well known that the transcriptional activity of structural anthocyanin biosynthesis genes is regulated by a ternary complex consisting of a MYB, a bHLH and a WD40 protein (MBW complex). The anthocyanin biosynthesis is even considered a model system for transcriptional control in eukaryotes. Previous studies identified transcriptional activation of different R2R3-MYBs[97–100] and bHLHs[101–103] as cause for the increase in anthocyanin levels. Our results showed that transcription factors were three times more often reported as causal factor of color differences than structural genes. While we cannot rule out the possibility that structural genes can also accumulate mutations prior to the reduction in transcription, this observation is in line with a previous study that observed faster evolutionary changes in transcription factors than in structural genes[95]. Similarly, Wheeler *et al*.[93] showed that transcription factors, particularly MYB, presented lower levels of gene expression with higher molecular evolutionary rate compared to their targeted structural genes, and suggested a negative correlation between evolution rate and gene expression in the Petunieae tribe[93]. This premise commonly known as the E-R anticorrelation, has been widly studied across different organisms, including yeast[104,105], Arabidopsis[106], Brassica[107], Barley[108], Arachis[109], and Drosophila[110]. However, the hypotheses explaining this model are still a topic of debate.

The loss of a transcription factor can switch off an entire biosynthesis pathway, while genes involved in this pathway could still be activated by other transcription factors to harness their activity in a different metabolic context. For example, *DFR* and *ANS*, two important anthocyanin biosynthesis genes, are also required for the biosynthesis of proanthocyanidins. A selective loss of anthocyanins and maintenance of the proanthocyanidin biosynthesis can thus not be caused by the loss of *DFR* or *ANS*. A well known example for such a scenario is the conservation of *DFR* and *ANS* across betalain-pigmented lineages of the Caryophyllales[111], which do not accumulate anthocyanins[96,112].

Studies on evolutionary rates investigating the components of the MBW complex suggested that MYBs would be the most likely component to be lost due to the highest degree of specialization which coincides with a lower pleiotropy[62,113,93]. It was observed that insertions/deletions were the most frequent mutation events in MYBs, while amino acid substitutions in the conserved region appeared irrelevant[61]. This let to the hypothesis that amino acid substitutions in transcription factors might not be relevant for the pigment evolution context, while InDels could disrupt the function of the encoded protein[61]. However, a recent investigation suggests a high importance of amino acid substitutions in the R3 interaction domain of MYBs in the loss of anthocyanin pigmentation in betalain-pigmented Caryophyllales[111]. These amino acid substitutions alter a highly conserved region that is considered crucial for the interaction of MYB and bHLH protein in the MBW complex which is required for activation of anthocyanin biosynthesis genes [13,114,115]. The lack of a functional MBW complex is considered as one crucial factor for the loss of anthocyanin pigmentation in the betalain-pigmented Caryophyllales[111]. In summary, there is evidence for InDels and amino acid substitutions as mechanisms that can disrupt the function of MYBs involved in the pigmentation biosynthesis regulation.

The frequency of mutations in TFs has been studied in various species. For example, a study of the *A. thaliana* genome revealed that there are more than 2,000 genes that encode for TFs and that these genes were more likely to accumulate mutations than non-TF genes[116]. Another study examined the frequency of point mutations in TF genes in *Eschericchia coli* and found that these genes were more vulnerable to harmful mutations that resulted in significant changes in gene expression than non-TF genes[117]. Given that their activity covers a wide range of functions, transcription factors can regulate and affect the expression of structural genes without the need for high gene expression levels[93]. A recent study by Liang *et al*. discovered a single point mutation in the 5’-UTR of *PELAN,* an anthocyanin-activating R2R3-MYB, as causal mutation for the loss of pigmentation in *Mimulus parishii*[118]. The results of their experiments concluded that despite the similar transcript abundance of *PELAN* in both strong pigmented and low pigmented cultivars, the difference in phenotype was due to a mistranslation of the *PELAN* mRNA[118].

The number of TFs required to regulate a specific enzyme-encoding gene is a complex and dynamic process that is influenced by various factors[119], including the complexity of the regulatory region[120], the stage of development or cell type[121], and environmental factors[122]. For example, only about ten *R2R3-MYBs* are known to bind specific DNA motifs related to the regulation of flavonoid biosynthesis pathway in *A. thaliana*[50,123]. In a review study, Feller *et al.*[58] performed a comparative analysis of the transcription factor families MYB and bHLH. They concluded that the reason why the bHLH family contains one of the largest numbers of transcription factors in plants is due to their functional diversification[58]. Multiple bHLH proteins contain a similar ligand-binding domain targeting different enzyme encoding genes[124]. A greater proportion of MYBs were recognized to be responsible for the regulation of flavonoid biosynthesis genes in comparison to bHLHs[124]. Our results align with this observation, because MYBs were reported as the causal gene of color differences in 75 cases, while bHLHs were only reported as crucial factor in ten cases. If bHLHs are more often involved in multiple processes, their loss would be more detrimental which makes it less likely to occur.

## CONCLUSION

A systematic literature screening supported the assumption that variations in transcription factors are the most frequently observed blocks in the anthocyanin accumulation in cases of intraspecies pigmentation differences. The MYB components of the MBW complex are dominating as blocks in the anthocyanin accumulation when comparing differently pigmented accessions. The degree of transcription factor specialization for a certain pathway seems to determine the frequency of their implication in color differences with more pleiotropic TFs like bHLH and TTG1 having a lower relevance. According to our results, MYBs are most often responsible for the difference in anthocyanin content, followed by bHLH, and other TFs. When structural genes appeared to be responsible for the absence of anthocyanins, this was most often a lack of DFR activity. An increased activation of the flavonol biosynthesis as a pathway competing for substrate with the anthocyanin biosynthesis was seen in rare cases.

## METHODS

### Extensive literature screening

An extensive record identification was performed in electronic databases (PubMed, Google Scholar, JSTOR) using specific screening keywords: “flower pigmentation differences”, “leaf pigmentation”, “color difference”, and “anthocyanin loss”. A total of 230 full-text articles were included for an eligibility assessment between December 2021 to October 2023. Accessible articles were considered if the genetic basis of pigmentation was investigated in the respective study (Additional file 1: Table S1). The evidence for causal genes reported in the literature differs between studies. We classified the presented evidences into the following main categories: ‘knockout mutant’, ‘in vitro characterization’, ‘coexpression patterns’, ‘metabolic exploration’, among others (Additional file 1: Table S1). Additionally, the respective family and order of each investigated species were collected along with the names of accessions, varieties, lines, or cultivars involved in the study. Most studies compared accessions of the same species that differed in anthocyanin pigmentation thus we are predominantly exploring intraspecific mechanisms of anthocyanin loss. While studies were classified as intraspecific or interspecific, we refrained from separate analyses due to a low sample size. As the classification of plants into categories like accessions, subspecies, and closely related species might leave some room for discussion, we considered all these studies.

### Data sources

From the 230 different articles, four studies were selected for in-depth transcriptome re-analysis. The four studies generated RNA-Seq data sets for the analysis of genetic factors underlying differences in pigmentation. The analyzed species were *Michelia maudiae*[125], *Rhododendron obtusum*[126]*, Trifolium repens*[127], and *Hosta plantaginea*[128] (Additional file 2). The selection of each dataset was based on the following criteria: (I) paired-end RNA-Seq data, (II) study has biological replicates, and (III) the authors have not identified the specific gene responsible for the color difference. The RNA-Seq datasets of those four plant species were retrieved from the Sequence Read Archive (www.ncbi.nlm.nih.gov/sra) (Additional file 1: Table S2) using fastq-dump[129].

### Transcriptome assembly

Transcriptomic data sets of four plant species were re-analysed to identify a candidate gene that could explain the absence of anthocyanins in one accession of each of these species. The generation of *de novo* transcriptome assemblies was necessary, because no transcriptome or genome sequences of these species were publicly available. Trimmomatic v0.39[130] was used to remove adapter sequences, to eliminate leading and trailing low-quality reads with quality below 3 (LEADING:3, TRAILING:3), and to drop reads shorter than 36 nt (MINLEN:36). The IDs of all remaining reads were modified by a customized Python script[131] to enable the following assembly with Trinity. Trinity v2.4.0[132] was applied for *de novo* transcriptome assembly using the previous cleaned reads as input. Trinity was run with a k-mer length of 25. In order to validate the quality of the transcriptome assemblies, a summary of the assembly statistics was generated for each species. The assembly statistics were computed using a previously developed Python script[133] (Additional file 1: Table S3).

After completion of the assembly process, kallisto v0.44[134] was run to quantify the abundances of transcripts based on all available RNA-Seq data of the respective species. Similarly, a principal components analysis (PCA) for every dataset was constructed based on their transcriptomic profiles to compare the similarity between samples and to identify any outliers (Additional file 3: Figures S1-S4). PCA was performed using R v.4.1.3[135] with the package ggplot2 v.3.4.0[136] to inspect the variation within the data set (Additional file 3: Figures S1-S4).

### Identification of Candidates Genes

In order to facilitate the identification of candidate genes, encoded peptide sequences were inferred from the transcriptome assembly using a previously established approach[137] that combines Transdecoder[138], ORFfinder[139], and ORFpredictor[140]. Knowledge-based Identification of Pathway Enzymes (KIPEs3) v0.34[141,142] was applied to identify the structural genes involved in the flavonoid biosynthesis. Flavonoid biosynthesis regulating R2R3-MYB genes were identified via MYB_annotator v0.3[143]. A previously described BLAST-based Python script[144] was deployed for the identification of additional candidates genes using bHLH, WD40, and WRKY genes associated with the flavonoid biosynthesis as baits[131]. A complete list of the selected candidate genes can be found in Additional file 1: Tables S4-S7.

Phylogenetic trees were constructed to identify all isoforms that belong to the same gene. Isoforms of the same gene may differ by the presence or absence of exons, but they should not exhibit more sequence differences than those accounted for by sequencing errors. In a phylogenetic context, transcript isoforms should form a distinct monophyletic group that can be replaced by one representative sequence. Phylogenetic trees were constructed with FastTree v.2.1.11[145] based on a MAFFT v7.475 alignment of polypeptide sequences using default parameters. Additional trees for comparison and additional support were constructed using IQ-TREE v.1.6.12[146] using default parameters and RAxML v.8.2.12[147] with PROTGAMMA+LG+F. Phylogenetic trees for the transcription factor families MYB, bHLH, TTG1, and WRKY were constructed to assess orthologues relationships (Additional file 2: Figures S8-S17). Sets of outgroup sequences were compiled based on reports in the literature in order to have a backbone of functionally characterized sequences for each tree. These sequences have been associated with the flavonoid biosynthesis in previous studies and were taken from plant species closely related to those explored with transcriptome assemblies.

### Taxonomic distribution of analyzed species

A plastid-based phylogenomic tree was used to study the distribution of the variations-related cases across all flowering plant families. The backbone tree was taken from Li *et al.*[77] and modified with iTOL v.6.8.1[148] to highlight the represented orders and families in our dataset.

### Gene expression analyses

Transcriptome assemblies usually generate a huge number of alternative transcript isoforms per gene. The initial sequences of the transcriptome assembly were used as reference for the quantification, but the transcript abundance (‘gene expression’) values were summarized per gene. Gene expression information of the entire monophyletic group was mapped to this representative transcript during the generation of heatmaps. Heatmaps displaying the candidate genes and their respective abundance as transcripts per million (TPM) were constructed using the R packages ComplexHeatmap v.2.10.0[149], circlize 0.4.15[150] and dplyr 1.1.0[151]. Genes with an adjusted p-value < 0.05 and absolute log2 fold-change > 1 were considered as differently expressed. The script used for the heatmap construction are available in our GitHub repository[131]. A workflow of the methods used in the transcriptome analysis is described in Figure 6.

**Figure 6.**
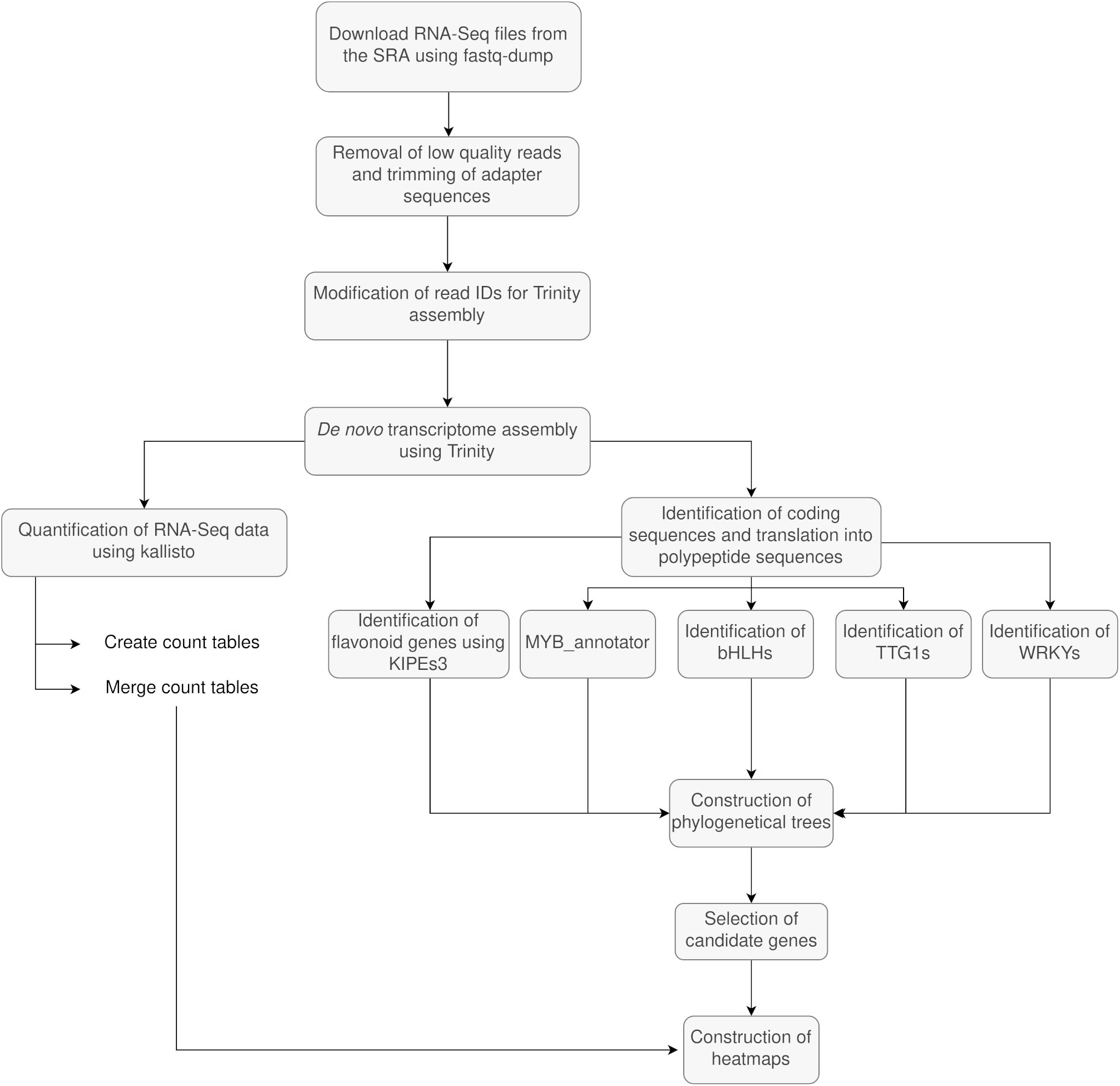
Flowchart representation of the methodology followed in the comparative transcriptional analysis. SRA – Sequence Read Archive, KIPEs – Knowledge-based Identification of Pathway Enzymes, MYB – Myeloblastosis, bHLH – basic helix-loop-helix, *TTG1* – *Transparent Testa Glabra 1*, WRKY – WRKY DNA-binding domain.

### Statistical Analysis

The observed distribution of frequency values in relation to the specific genes hotspots of color differences in the 230 studies, were evaluated using a 2-factor Chi-Square test. All the tests were applied using the statistical software R v.4.1.3[135].

## Abbreviations

PAL: phenylalanine ammonia-lyase
C4H: cinnamic acid 4-hydroxylase
4CL: 4-coumarate-CoA ligase
CHS: chalcone synthase
CHI: chalcone isomerase
F3H: flavanone 3-hydroxylase
F3’H: flavonoid 3’-hydroxylase
F3’5’H: flavonoid 3’,5’-hydroxylase
DFR: dihydroflavonol 4-reductase
ANS: anthocyanidin synthase
UFGT: UDP-glucose:flavonoid 3-O-glucosyltransferase
FLS: flavonol synthase
ANR: anthocyanidin reductase
LDOX: leucoanthocyanidin dioxygenase
FNS: flavone synthase
ER: Endoplasmic Reticulum
MATE: Multidrug and Toxin Extrusion Transporter
GST: Glutathione S-transferase
ABCC: ATP-binding cassette
SRA: Sequence Read Archive
KIPEs: Knowledge-based Identification of Pathway Enzymes
MYB: Myeloblastosis
bHLH: basic helix-loop-helix
TTG1: Transparent Testa Glabra 1
WRKY: WRKY DNA-binding domain.

## Acknowledgements

This work was supported by the BMBF-funded de.NBI Cloud within the German Network for Bioinformatics Infrastructure (de.NBI) (031A532B, 031A533A, 031A533B, 031A534A, 031A535A, 031A537A, 031A537B, 031A537C, 031A537D, 031A538A). We are grateful to de.NBI and the Center for Biotechnology (CeBiTec) at Bielefeld University for providing us with the resources and setting to conduct the necessary computational analyses. We are also thankful to the members of the Plant Biotechnology and Bioinformatics group at TU Braunschweig for their helpful discussions and valuable comments during the manuscript revision process. We acknowledge support by the Open Access Publication Funds of Technische Universität Braunschweig. We used bioRender.com for the construction of some figures.

## Declarations

### Authors’ contributions

BP conceptualized the research project. MMR carried out the analyses and prepared figures. MMR and BP wrote the manuscript. Both authors read and approved the final manuscript.

## Funding

This work received no external funding.

### Availability of data and materials

All the transcriptome data used in this study was obtained from the NCBI Sequence Read Archive (SRA) under accession numbers PRJNA504531, PRJNA542483, PRJNA700000, and PRJNA393638 (http://www.ncbi.nlm.nih.gov/sra). The data charts supporting the results and conclusions are included in the article and additional files. All the assemblies and scripts used in reanalyzed transcriptome analysis have been deposited in our GitHub repository (https://github.com/bpucker/codi).

### Ethics approval and consent to participate

Not applicable.

### Consent for publication

Not applicable.

