## Additional file 2 for "Genetic factors explaining anthocyanin pigmentation differences"

*Michelia maudiae*

A study about the flower color variation in *Michelia maudiae*[1] generated RNA-Seq data sets (Additional file 1: Table S2) of plants of an anthocyanin-pigmented accession (red flower) and of plants of an unpigmented accession (white flower). Flower samples were taken at an early and a late developmental stage to compare not only accessions, but also developmental stages. We identified the transcript sequences of flavonoid biosynthesis genes in our transcriptome assembly of *M. maudiae*[2]. The results from our re-analyses showed high expression levels of *F3H* and *TT19* in the red flowers regardless of the development stage. The gene *ANR* showed high expression in the white flowers particularly in the late developmental stage. The gene expression of *DFR* had a slightly increased expression in the red late flowers (Figure 1A).

The genes related to the WD40 transcription factor TTG1 (Figure 1B) presented higher expression values in the red flowers independently of the developmental stage. Gene expression of *TTG1* in the red flower was considerably higher at the late developmental stage compared to the early one. Members of the transcription factor families R2R3-MYB, bHLH, WD40 and WRKY were also identified and analyzed with respect to their gene expression. Three genes related to the R2R3-MYB transcription factors associated with the flavonoid biosynthesis were detected. From those, the repressor *MYB4* showed a high expression at the late stage of red flowers. In addition, a slight increase in expression over the different developmental stages was observed for the genes *MYB12, MYB11, MYB111,* and *MYB75, MYB90, MYB113, MYB114* (Figure 1C). Two genes of the bHLH transcription factor family associated with the regulation of the flavonoid biosynthesis were discovered. One gene is related to Lily Hybrid bHLH1 (*LhbHLH1)* and the other one to *TT8* (Figure 1D). The expression of the genes related to *TT8* were not particularly different between the flower stages. On the contrary, the genes related to Lily Hybrid bHLH1 (*LhbHLH1)* showed a higher expression in samples taken from the red flowering accession at the early stage.  Furthermore, in the transcription family WRKY (Figure 1E) four different genes were identified as putative orthologs of the anthocyanin regulators reported in *Malus domestica and Pyrus spp.* Orthologs of the genes MdWRKY40, and PyWRKY31 presented the higher expression in samples of the red flowers compared to the white flowers at the early developmental stage. In summary, there are several differences in gene expression like the strong expression of *TT19* in the red samples and the almost complete absence of expression of this gene in the white samples which suggest that this step could be the bottleneck in the anthocyanin accumulation which ultimately results in white flowers. However, the gene expression differences of other genes in the anthocyanin biosynthesis suggest that the causal gene is more likely to be a transcription factor that activates multiple structural genes.

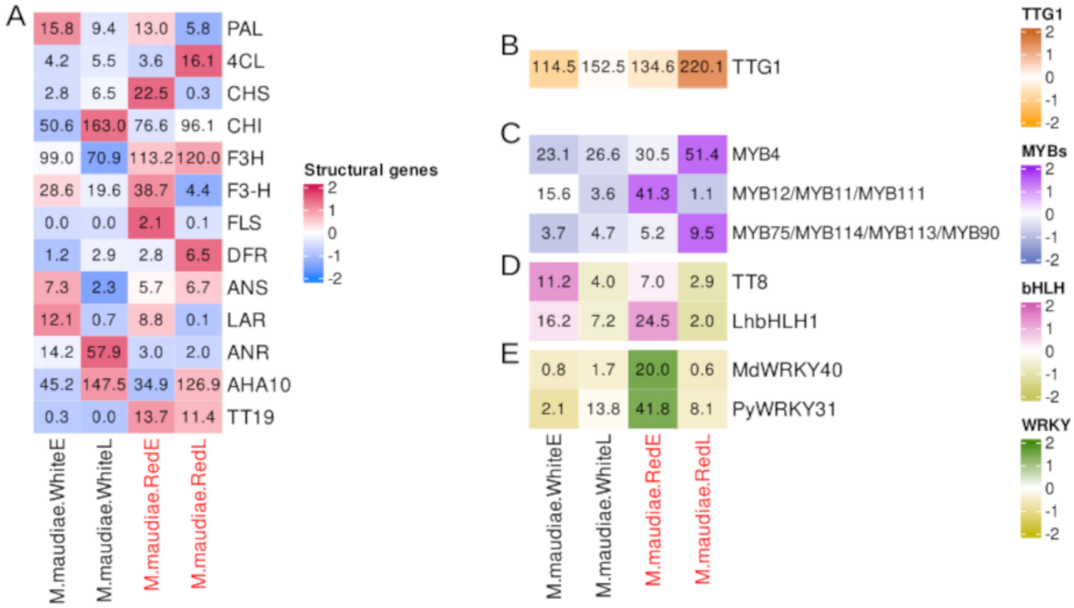


**Figure 1.** Heatmaps presenting gene expression values of white and red *Michelia maudiae* flowers and their respective developmental stages, early (E) and late (L). (A) Structural genes, (B) homologs of *TTG1*, (C) homologs of MYBs, (D) homologs of bHLH, and (E) homologs of WRKY. An isoform resolved version of this heatmap is available in Additional file 3: Figure S5.

*M. maudiae* is a perennial tree belonging to the family Magnoliaceae. Lang *et al.* [1] conducted an RNA-Seq experiment to investigate the flower color variation in *M. maudiae.* Early and late developmental stages of the white and red flowers were sampled. The original analysis did not reveal an individual gene responsible for the pigmentation difference. Our reanalysis revealed high expression of the known anthocyanin transporter gene *TT19 (GST)* [3] in the red flowers while there was no substantial gene expression in the white flowers. This suggests that a variation in the promoter of the *TT19* ortholog or its regulators could prevent the anthocyanin accumulation in the white flowering cultivar. Variations disrupting this gene were previously reported as blocks in the anthocyanin accumulation [4–6]. An established hypothesis states that this protein protects anthocyanins during transport to the central vacuole [6]. In addition, the structural genes *CHI, CHS, F3H, DFR and ANR* presented the highest expression levels in the red flowers compared to their expression in the white ones. It is unlikely that five structural genes would be up-regulated independently and more likely that a transcription factor is the causal factor in this scenario. Furthermore, we identified two transcription factor genes, one ortholog related to the repressor *AtMYB4* [7], and one ortholog of *LhbHLH1* known to be involved in the anthocyanin biosynthesis in *Lilium spp.* [8]. Clear differences in the expression of unigenes related to the bHLH transcription factor were determined. In red flowers, genes related to *LhbHLH1* showed higher expression levels compared to white flowers. Additionally, the higher expression of WRKY transcription factor-related genes in red 'Rubellis' flowers compared to white flowers is of particular interest. Similar results were reported by Liu *et al*. [9] The authors reported up-regulation of bHLH and WRKY transcription factor-related genes only at the early stage of flowers belonging to the species *Michelia crassipes*. According to their claims, the reason why bHLH and WRKY genes appear to be up-regulated in the early stages of flower development could be due to their importance in petal color transition during flowering. Recent studies hypothesize that the regulation of the production of anthocyanin pigments is not entirely handled by the MBW complex, but that transcription factors such as those belonging to the WRKY family may also play a role in the regulation and transport of these pigments [10–12].

*Rhododendrom obtusum*

A study about the flower coloration of *Rhododendron obtusum* [13] generated a series of RNA-Seq datasets that cover five developmental stages of two varieties. The variety “Yanzhi Mi” characterized by a bright pink color and the wild type variety “Dayuanyangjin” which presents white petals with pink stripes. The authors of this study state that derivatives of cyanidin, peonidin, and pelargonidin might be responsible for the pink color of mutant petals and that the D2 stage was the key stage of flower color formation. Since no reference sequence is publicly available, a *de novo* transcriptome assembly of *R. obtusum* was generated for the identification of transcripts belonging to flavonoid biosynthesis genes [2].

Results of the re-analyses showed high expression values in stage two (D2) of the white variety in genes *AHA10, UGT72L1, ANR,* and *A3GT* in comparison to the pink flowers at that same stage. On the contrary, the pink flowers showed high expression for the genes *4CL, LAR,* and *MATE* at stage two (Y2). No other particular difference between the two varieties was observed in the other stages with the exception of *F3'5'H* (Figure 2A).

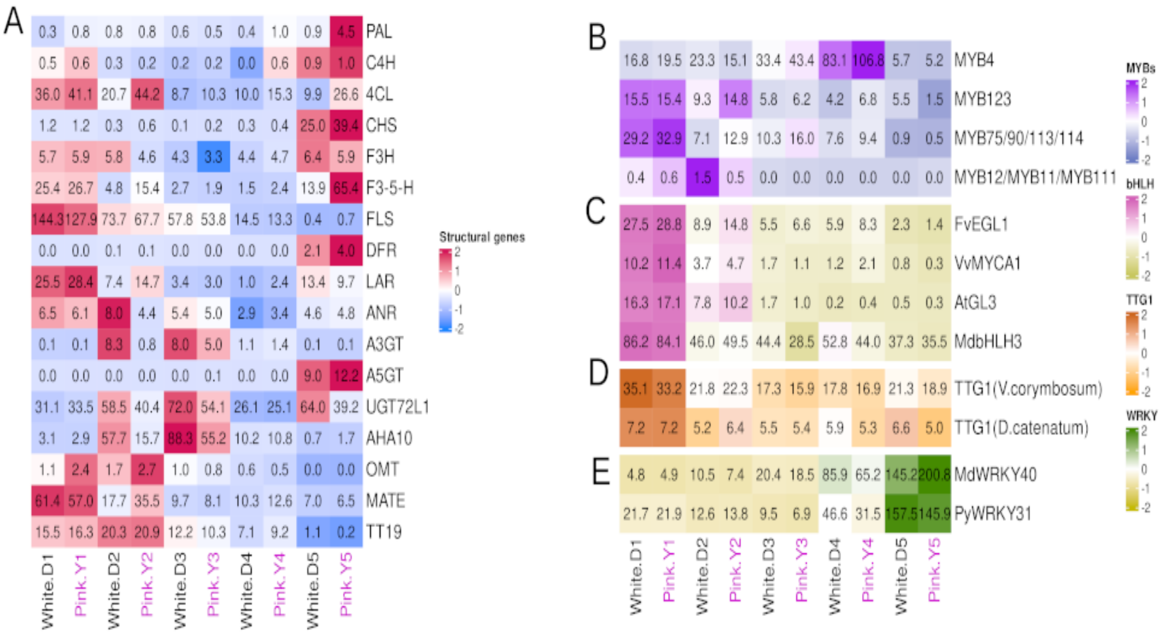

**Figure 2.** Heatmaps presenting gene expression values of white flowering *Rhododendron obtusum* var. *'Dayuanyangjin'* in different developmental stages: closed buds (D1), buds showing color at the top (D2), the initial flowering stage (D3), the full flowering stage (D4) and the last flowering stage (D5)  and pink flowering *Rhododendrom obtusum* var. *'Yanzhi Mi'* with the different flower stages: closed buds (Y1), buds showing color at the top (Y2), the initial flowering stage (Y3 stage), the full flowering stage (Y4 stage) and the last flowering stage (Y5) (A) Structural genes, (B) homologs of *TTG1*, (C) homologs of MYBs, (D) homologs of bHLH, and (E) homologs of WRKY. An isoform resolved version of this heatmap is available in Additional file 3: Figure S6.

Two *TTG1* genes belonging to two different clades of the species *Vaccinium corymbosum* and *Dendrobium catenatum* were identified. These genes did not show a clear difference in gene expression between the pink and white samples at any developmental stage (Figure 2B). Three genes related to the *A. thaliana* transcription factors *AtMYB75, AtMYB123,* and *AtMYB4* (Figure 2C) were identified. According to Sun *et al*. [13], D2 is the developmental stage where the difference in pigmentation becomes visible. This analysis revealed a difference in expression of the structural genes in D2 samples. It is also interesting to notice a slight difference in the expression of genes related to the transcription factor families MYB and bHLH between the same two D2 groups. This observation suggests a possible involvement of MYB and bHLH in the regulation divergence (Figure 2D). The WRKY transcription factor family shows no particular difference in expression between the varieties at any developmental stage (Figure 2E). In summary, it seems that a slightly higher activity of *MYB75* and *FvEGL1/AtGL3* orthologs could be responsible for the strong pigmentation of the pink cultivar.

*R. obtusum* commonly known as Japanese azaleas is a small shrub, characterized by its pink flowers. Sun *et al*. [13] provided a starting point into the genetic mechanisms underlying the flower color divergence in the *R. obtusum* species by generating an RNA-Seq data set. They analyzed the gene expression in two differently pigmented varieties of *R. obtusum* in five flower developmental stages. The variety "Yanzhi Mi" characterized for a bright pink color and the wild type variety "Dayuanyangjin" which presents white petals with pink stripes. Their results indicated a high gene expression in *CHS, CHI, F3Hs,* and *F3'H* in the pink variety in stage 2 of flower development [13]. According to the authors, developmental stage two was considered as the key point of divergence in flower pigmentation between 'Yanzhi Mi' variety (pink flowers) and the 'Dayuanyangjin' variety (white flowers). Our results showed a slight difference in gene expression mainly of *FLS* at stage two of white flowers (D2), which could indicate a channeling of substrate towards flavonol production. High activity of the flavonol biosynthesis is consistent with the absence of red anthocyanins in the flowers as previously reported in numerous species [14–16]. Moreover, an increased expression of the transporter gene *MATE* (*TT12*) in the pink flowers could explain the anthocyanin accumulation starting at the stage 2 of flower development [17]. With respect to the variety 'Yanzhi Mi' an increase in the expression of unigenes related to *MYB75* and *GL3* of the bHLH family was identified. Our findings support the results of a recent study on *Rhododendron pulchrum* species [18], which presented a comparative transcriptome analysis between three varieties with different phenotypes in flower color. Their results also determined a high expression of a gene encoding the FLS enzyme, increasing the flavonol content at the expense of anthocyanin formation in the variety with white flowers. At the same time, this variety showed increased expression of *MYB5*, which they claimed negatively regulates anthocyanin synthesis [18].

*Trifolium repens*

*Trifolium repens* commonly known as "white clover" is a perennial herbaceous plant native to Europe and western Asia. This plant is characterized by its white flower, however plants with red or pinkish flowers can be seen in rare cases. Zhang *et al.* [19], observed that those red flowering clover varieties produce white flowers under shaded conditions. Therefore, they performed an RNA-Seq analysis on the anthocyanin biosynthesis of a red clover variety to identify the genes associated with the flower color. In their results eight candidates genes were identified. Most of them were down-regulated in plants under shaded conditions (*CHS, F3'H, F3'5'H, ANS, UFGT, and DFR*) and only two genes were found to be up-regulated (*LAR* and *FLS*) [19].

A *de novo* transcriptome assembly of *T. repens* was generated for the identification of transcripts belonging to flavonoid biosynthesis genes [2]. The results of our transcriptome re-analysis (Figure 3A) confirmed the low expression levels of the structural genes *CHS, F3'H, F3'5'H, DFR, ANS, TT19, and UFGT,* for the white flowers in comparison with the red flowers. However, these white flowers presented relatively high expression of the genes *4CL*, *AHA10*, and *TT15*. Additionally, we analyzed the expression levels of members of the MYB, bHLH, *TTG1,* and WRKY transcription factor families.

One ortholog of the *A. thaliana MYB75* was identified (Figure 3B). No distinct pattern between the varieties was observed suggesting MYBs are not likely to be responsible for the color difference. By contrast, the family bHLH (Figure 3C) presented differences between the red and white samples, particularly the unigenes related to the transcription factors *Lonicera japonica GL3* and *Arabidopsis EGL3*. This implies that a possible inactivation of bHLH due to lack of light exposure could be associated with the loss of pigmentation in the white flowers of *T. repens*. Concerning the transcription factor *TTG1* (Figure 3D) no substantial difference in expression was identified between the two groups. In the WRKY transcription factor family (Figure 3E) the red flowers presented higher expression levels, especially the genes related to *M. domestica WRKY40* and *Pyrus spp. WRKY31*. In summary, higher bHLH expression in light exposed clover plants, especially of the encoding genes of TFs *EGL3* and *GL3,* appears to be the most likely explanation for the generally increased expression levels of various structural genes.

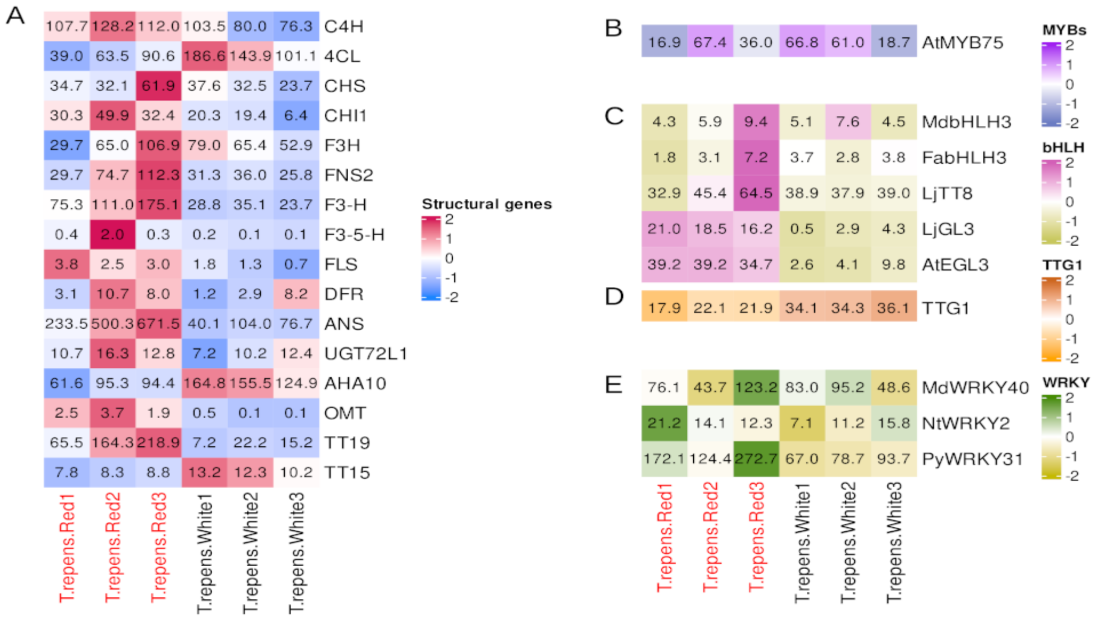


**Figure 3.** Heatmaps presenting gene expression values of red flowering Trifolium repens and white flowering *T. repens* exposed to shaded conditions. (A) Structural genes, (B) homologs of MYBs (C) homologs of bHLHs, (D) homologs of TTG1, and (E) homologs of WRKYs. An isoform resolved version of this heatmap is available in Additional file 3: Figure S7.

Trifolium is a worldwide spread genus belonging to the Fabaceae family. Adaptation to different environments has made this herbaceous plant to present a great variety of phenotypes, especially variations in the color pattern of its leaves and flowers [20]. The study by Zhang *et al.* [19] concluded that low expression of *CHS* and *F3'H* in the shaded plants likely limits the formation of dihydrokaempferol, which results in a blockage in the production of the red-magenta cyanidin pigment.

Similarly, our results revealed an increase in expression of the F3’H encoding gene in the red flowers. This upregulation of *F3’H* might suggest a redirection of substrate towards dihydroquercetin, which promotes the production of anthocyanins, specifically of the cyanidin class that results in the red coloration. High expression of *ANS* is supporting this hypothesis. Despite being a critical component in the substrate contribution of the flavonoid biosynthetic pathway, the F3'H enzyme is frequently overlooked in discussions of DFR and FLS specificity. For instance, the expression of *F3'H* in grapevines increases when grapes turn red during veraison[21]. Similar results were observed in *Fragaria vesca* and *Vaccinium corymbosum* with an increase of *F3'H* transcript levels during fruit ripening [22,23]. Additionally, a clear difference in bHLH expression was observed, especially in *EGL3* and *GL3*, between flowers exposed to light (red flowers) and those kept in shade (white flowers). Flowers grown under shaded conditions showed a low expression of these bHLHs. Similar results were presented in a study by Hong *et al*. [24] They performed a transcriptomic analysis in *Chrysanthemum* to identify the effects of light intensity on anthocyanin synthesis. The results presented by Hong *et al*. showed changes in the coloration of ray florets when their capitula were exposed to different shading conditions. Those capitula that were exposed to normal light developed into pink flowers while those that were exposed to dark conditions developed into white or pale pigmented flowers, demonstrating the effect that light has on the accumulation of anthocyanins in plant tissues. However, the results of transcriptomic analysis also identified that the genes encoding the bHLH protein showed low gene expression in those plants incubated in the shade.

Research involving different light conditions and anthocyanin synthesis has been a subject of discussion for many years. Early studies proposed that the transcription factor bZIP, together with the MBW complex, would regulate the anthocyanin formation in different plant tissues in response to light conditions [25,26]. Now, it is known that the molecular mechanism behind the impact of shade on plant color production is more complex and depends on the type of plant and the duration of the light exposure [27]. Other important factors are phytohormones and HY5, a TF of the bZIP family, which positively regulates the expression of genes related to anthocyanin synthesis upon light exposure [28,29]. The evident inactivity of bHLH in the flowers of plants growing in the shade might be responsible for the lack of *F3'H* activity that led to the absence of pigmentation in the flowers of *T. repens.* Similar results have been observed in purple broccoli (*Brassica oleraceae*) when cultivated in the shade [30].

*Hosta plantaginea*

A study was conducted to investigate anthocyanin accumulation in the flowers of the herbaceous plant Hosta plantaginea[31]. The study generated RNA-seq datasets for three different cultivars: *Hosta plantaginea* plantaginea (HP), Hosta Cathayana (HC), and Hosta 'Summer Fragrance' (HS). Each cultivar exhibited different flower pigmentation. Cultivar HP had white flowers, cultivar HC had flowers with a subtle purple coloration, and cultivar HS had dark purple flowers. Additionally, three developmental stages were analyzed for each cultivar: flower bud stage (HP-1, HC-1, and HS-1), initial flowering stage (HP-2, HC-2, and HS-2), and late flowering stage (HP-3, HC-3, and HS-3).

The initial study reported increased expression levels of ANS and CHS during the late flowering stage compared to the bud and initial stages [31]. However, the specific regulatory genes responsible for the differences in color pigmentation between the three cultivars were not identified. Our results demonstrate high expression levels of CHS, DFR, and ANS in the late stages of HC and HS, respectively (Figure 4A). Moreover, F3H and FLS exhibit high expression levels in all stages of the light purple HC and white HP, suggesting an increase in the production of colorless flavonols. Particularly interesting is the high expression of F3'5'H observed only in cultivar HC, while LAR, but not ANR, shows high expression exclusively in different stages of HP flowers.

Regarding the regulatory genes, we identified the repressor MYB4 and an ortholog related to the genes AtMYB75, AtMYB90, AtMYB113, or AtMYB114 (Figure 4C). Among these, high expression was only observed in the late stage of HC, with no significant difference between the HP and HS varieties. The repressor MYB4 showed high expression in the late stage of HP, with no particular difference between the HC and HS varieties. Additionally, two orthologs related to the genes bHLH1 and bHLH2 were identified. Higher expression of bHLH2 was observed in the late stages of all three varieties. Similar tendencies were observed for bHLH1, except that HP exhibited very low expression in the late stage (Figure 4D).

Two clades related to genes belonging to the WD40 family (TTG1) were identified (Supplementary file 2: Figure S17A). In these clades, four genes and two genes were found, respectively. However, there was no difference in gene expression between the three varieties and developmental stages (Figure 4B). Interestingly, higher expression levels were observed for the transcription family WRKY in the HC variety compared to the HP and HS varieties (Figure 4E).

In summary, the high activity of FLS in white and less pigmented flowers indicates a shift in substrate utilization towards the production of flavonols, leading to white flowers or flowers with lighter colors. This premise is directly supported by the observed absence of DFR and ANS expression in the white cultivar.

Although our analysis provides a potential explanation for the variation in pigmentation among the cultivars, we have chosen not to include this species in the main results section due to inconclusive findings and the inability to pinpoint the specific gene responsible for the observed differences in pigmentation between the three cultivars.

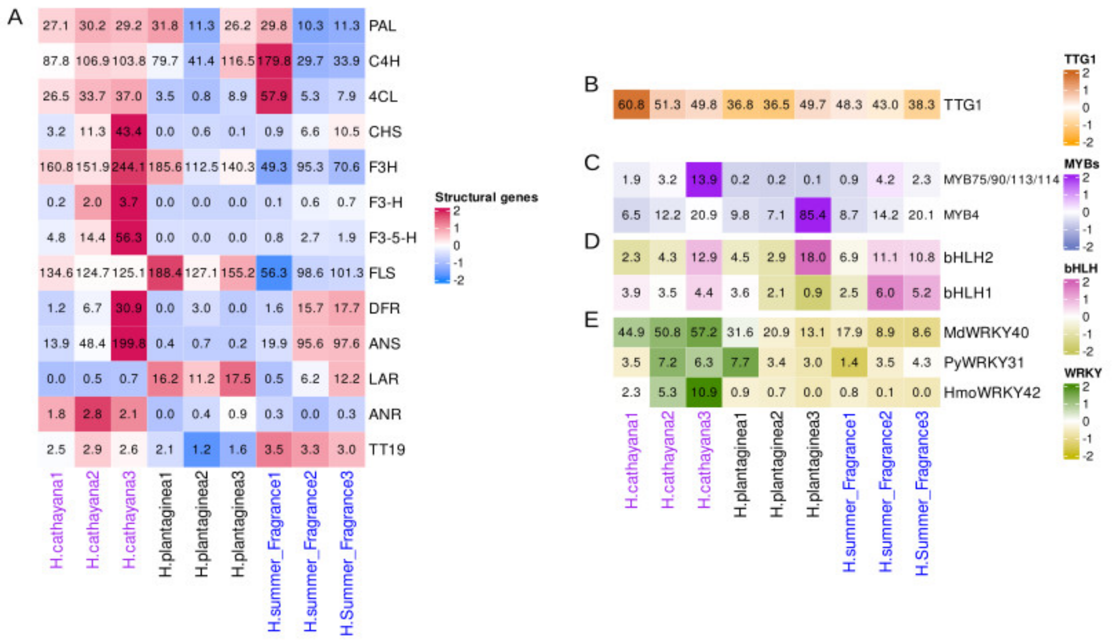


**Figure 4.** Heatmaps presenting gene expression values of *Hosta plantaginea* var. 'cathayana' (HC) with a light lila color, *Hosta plantaginea* var. 'plantaginea' (HP) with white flowers represented with the black text, and *Hosta plantaginea* var. 'summer' (HS) which presented deep purple flowers and is shown in this figure with blue colored text. Each variety is shown in different developmental flower stages (1, 2, and 3). (A) Structural genes, (B) homologs of TTG1, (C) homologs of MYBs, (D) homologs of bHLHs, and (E) homologs of WRKYs.

10. Lloyd A, Brockman A, Aguirre L, Campbell A, Bean A, Cantero A, et al. Advances in the MYB–bHLH–WD

Repeat (MBW) Pigment Regulatory Model: Addition of a WRKY Factor and Co-option of an Anthocyanin MYB for Betalain Regulation. Plant Cell Physiol. 2017 Sep 1;58(9):1431–41.

27. Araguirang GE, Richter AS. Activation of anthocyanin biosynthesis in high light - what is the initial signal?

New Phytol. 2022 Dec;236(6):2037–43.

28. Ma Y, Ma X, Gao X, Wu W, Zhou B. Light Induced Regulation Pathway of Anthocyanin Biosynthesis in Plants. Int J Mol Sci. 2021 Oct 15;22(20):11116.

29. Xie G, Zou X, Liang Z, Wu D, He J, Xie K, et al. Integrated metabolomic and transcriptomic analyses reveal molecular response of anthocyanins biosynthesis in perilla to light intensity. Front Plant Sci. 2022;13.

30. Liu C, Yao X, Li G, Huang L, Xie Z. Transcriptomic profiling of purple broccoli reveals light-induced anthocyanin biosynthetic signaling and structural genes. PeerJ. 2020 May 5;8:e8870.

31. Zhang J, Sui C, Wang Y, Liu S, Liu H, Zhang Z, et al. Transcriptome-Wide Analysis Reveals Key DEGs in Flower Color Regulation of *Hosta plantaginea* (Lam.) Aschers. Genes. 2020 Jan;11(1):31.
