## Additional file 3 for "Genetic factors explaining anthocyanin pigmentation differences"

### SUPPLEMENTARY FIGURES

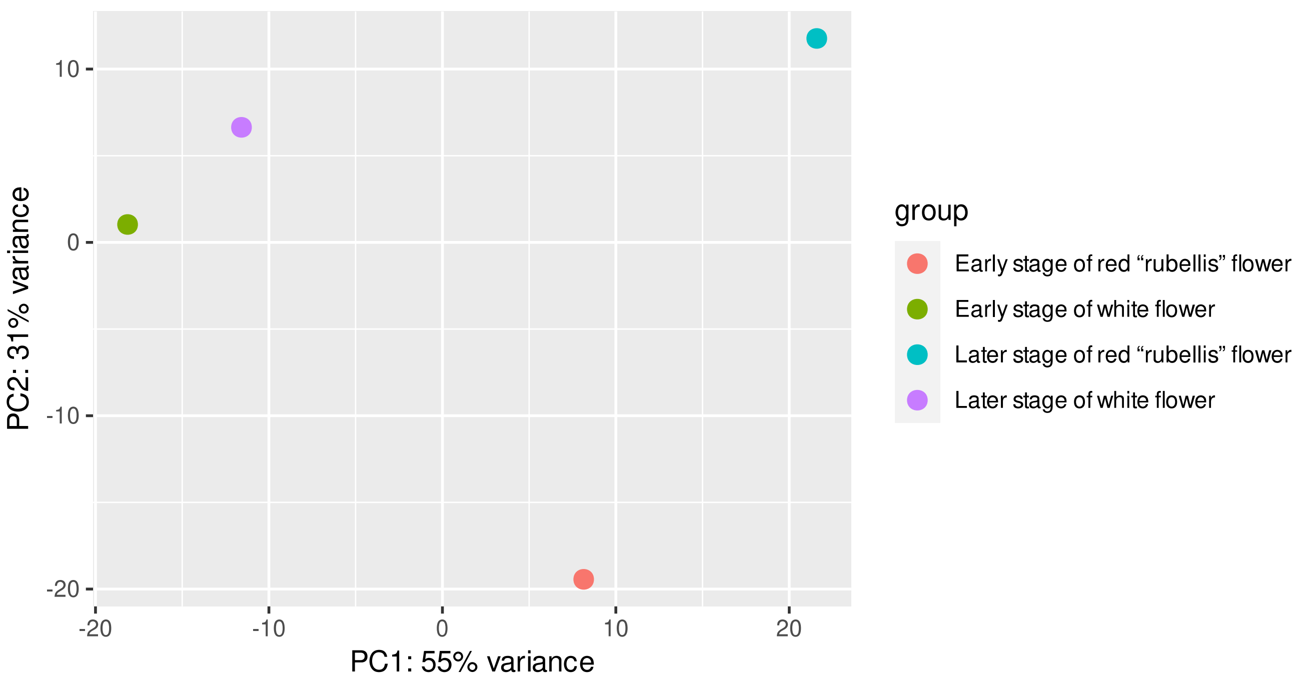


**Figure S1.** Principal Components Analysis *Michelia maudiae*

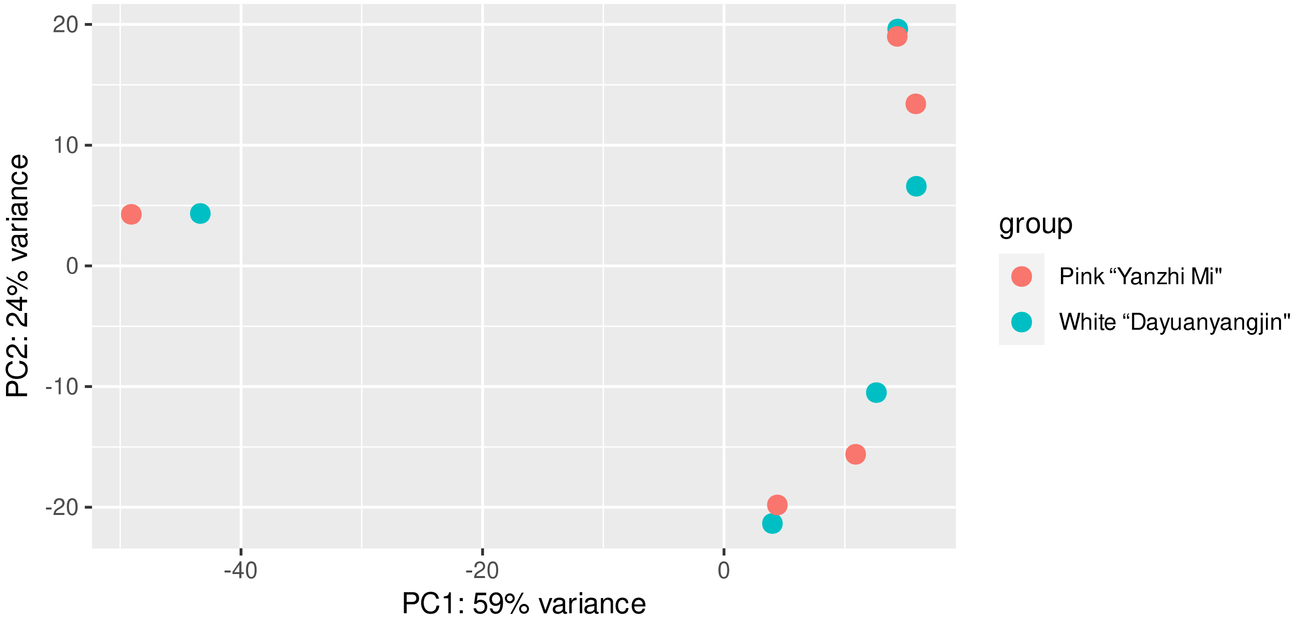


**Figure S2.** Principal Components Analysis *Rhododendron obtusum*

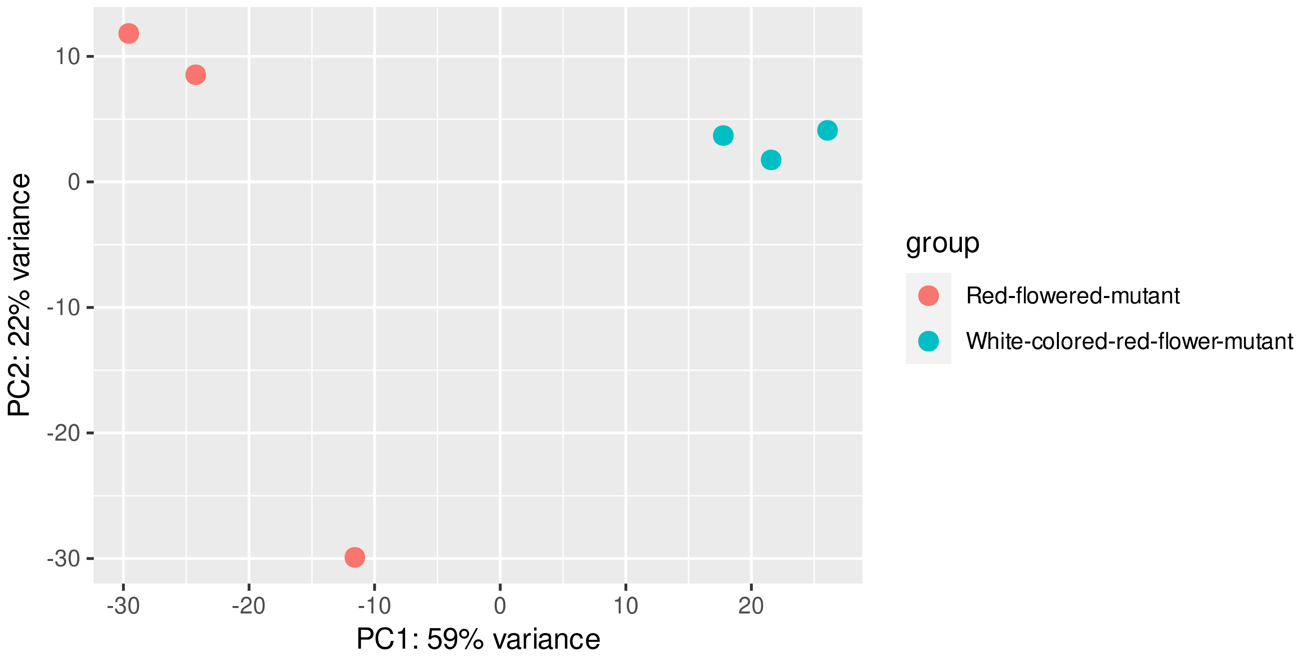


**Figure S3.** Principal Components Analysis *Trifolium repens*

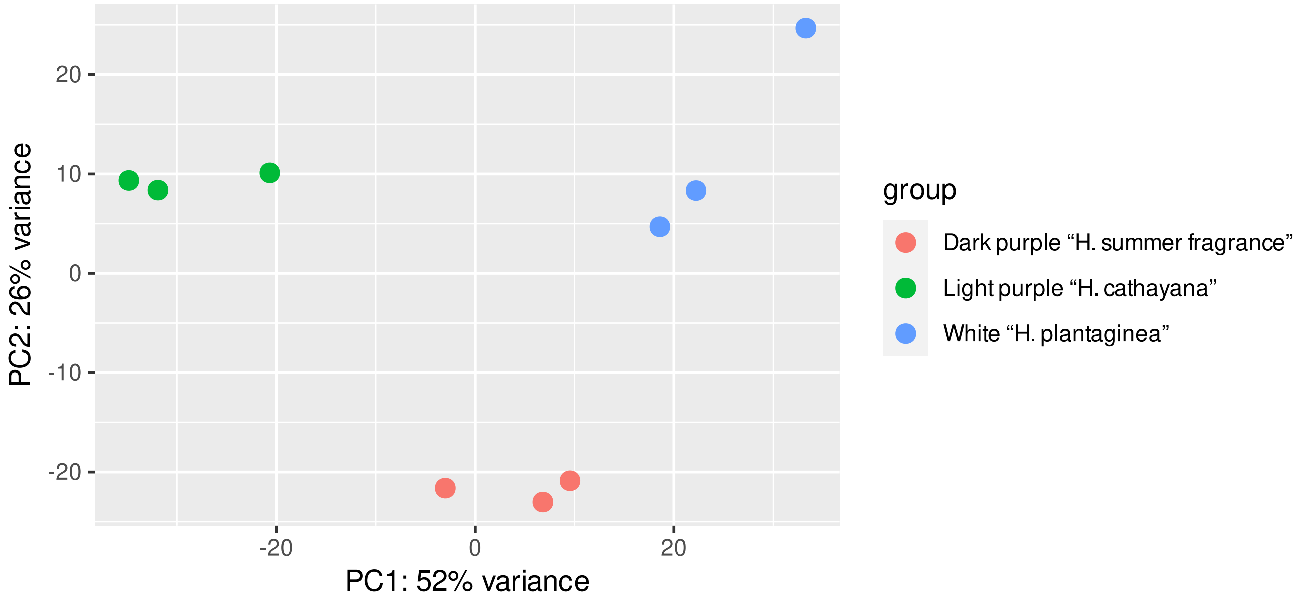


**Figure S4.** Principal Components Analysis *Hosta Plantaginea*

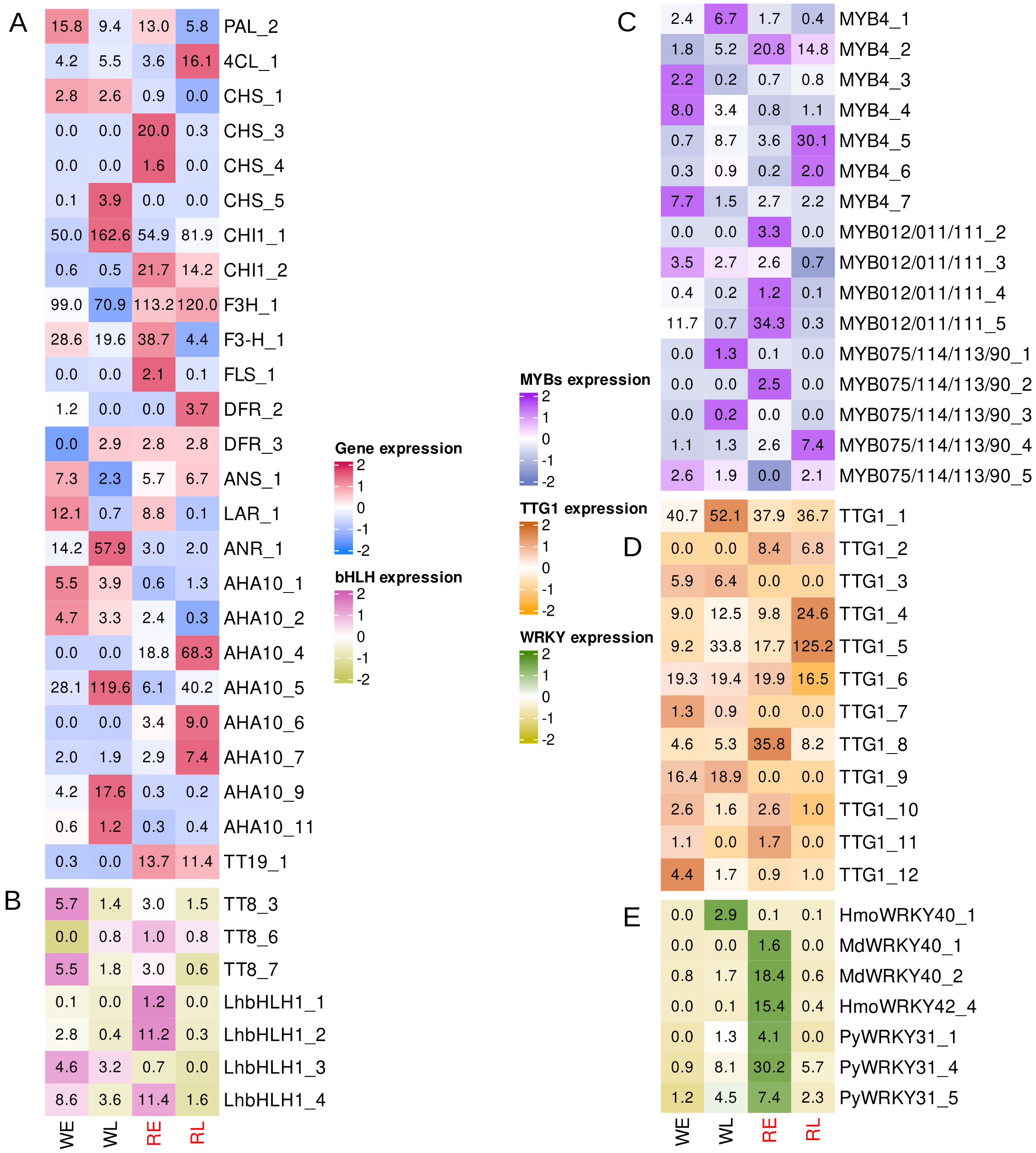


**Figure S5.** Complete isoform heatmaps presenting gene expression values of white (W) and red (R) *Michelia maudiae* flowers and their respective developmental stages, early (E) and late (L). (A) Structural genes, (B) bHLHs, (C) MYBs, (D) TTG1, and (E) WRKY.

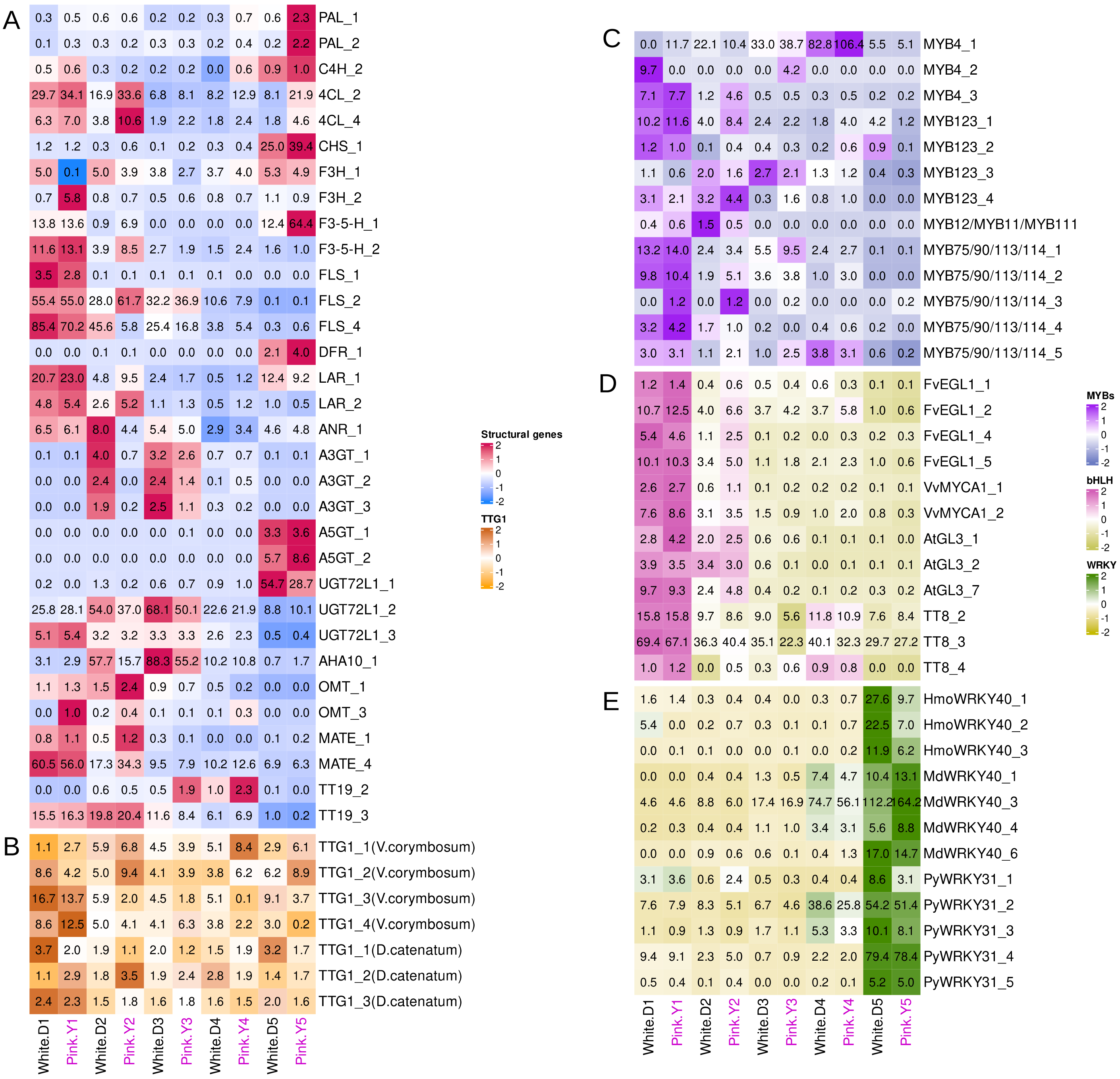


**Figure S6.** Complete isoform heatmaps presenting gene expression values of white flowering *Rhododendron obtusum* var. *'Dayuanyangjin'* in different developmental stages: closed buds (D1), buds showing color at the top (D2), the initial flowering stage (D3 stage), the full flowering stage (D4 stage) and the last flowering stage (D5) and pink flowering *Rhododendrom obtusum* var. *'Yanzhi Mi'* with the different flower stages: closed buds (Y1), buds showing color at the top (Y2), the initial flowering stage (Y3 stage), the full flowering stage (Y4 stage) and the last flowering stage (Y5) (A) Structural genes, (B) MYBs, (C)bHLHs, (D) TTG1, and (E) WRKYs.

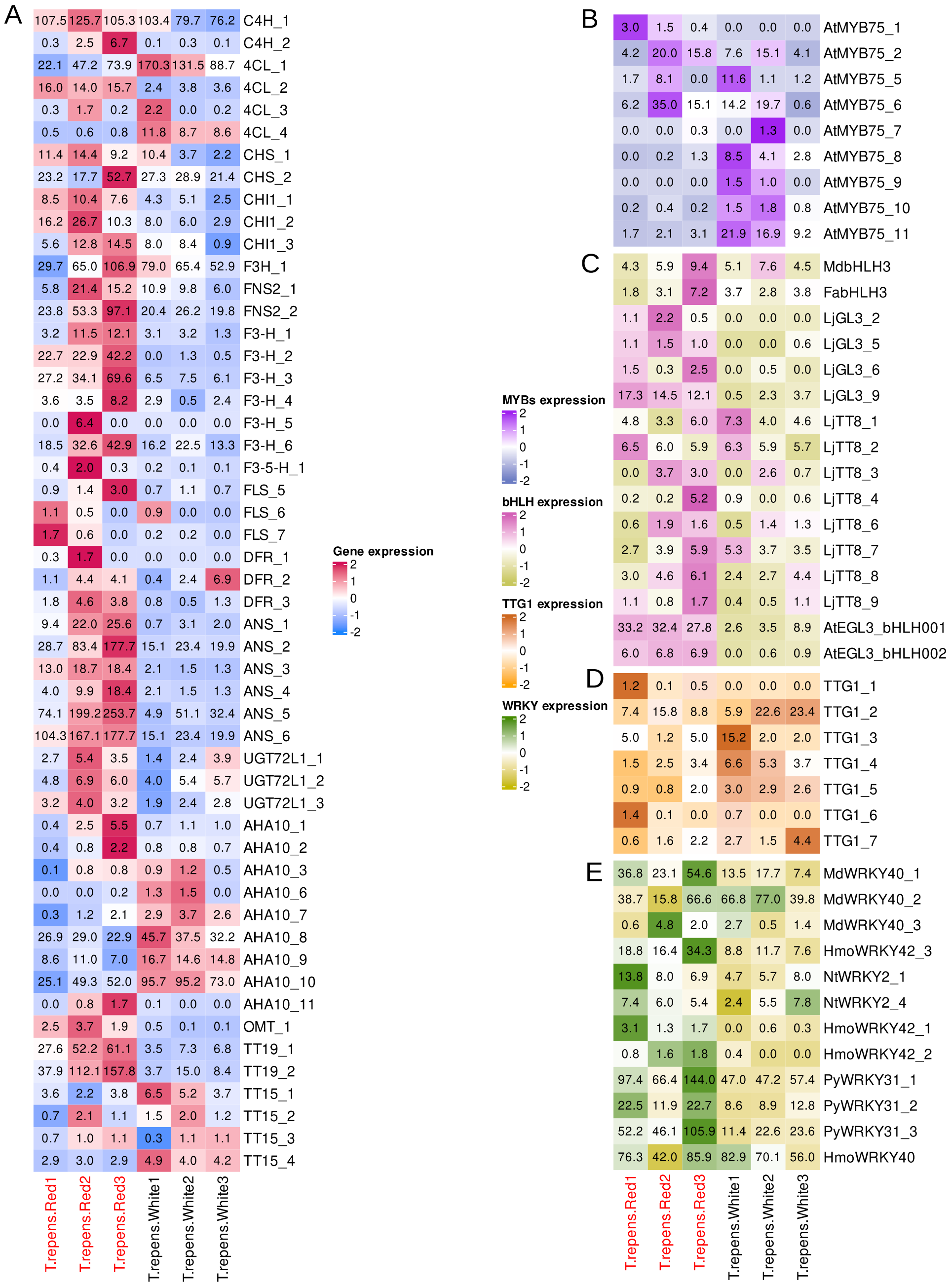


**Figure S7.** Complete isoform heatmaps presenting gene expression values of red flowering *Trifolium repens* and white flowering *T. repens exposed to shaded conditions* (A) Structural genes, (B) MYBs, (C) bHLHs, (D) TTG1, and (E) WRKYs

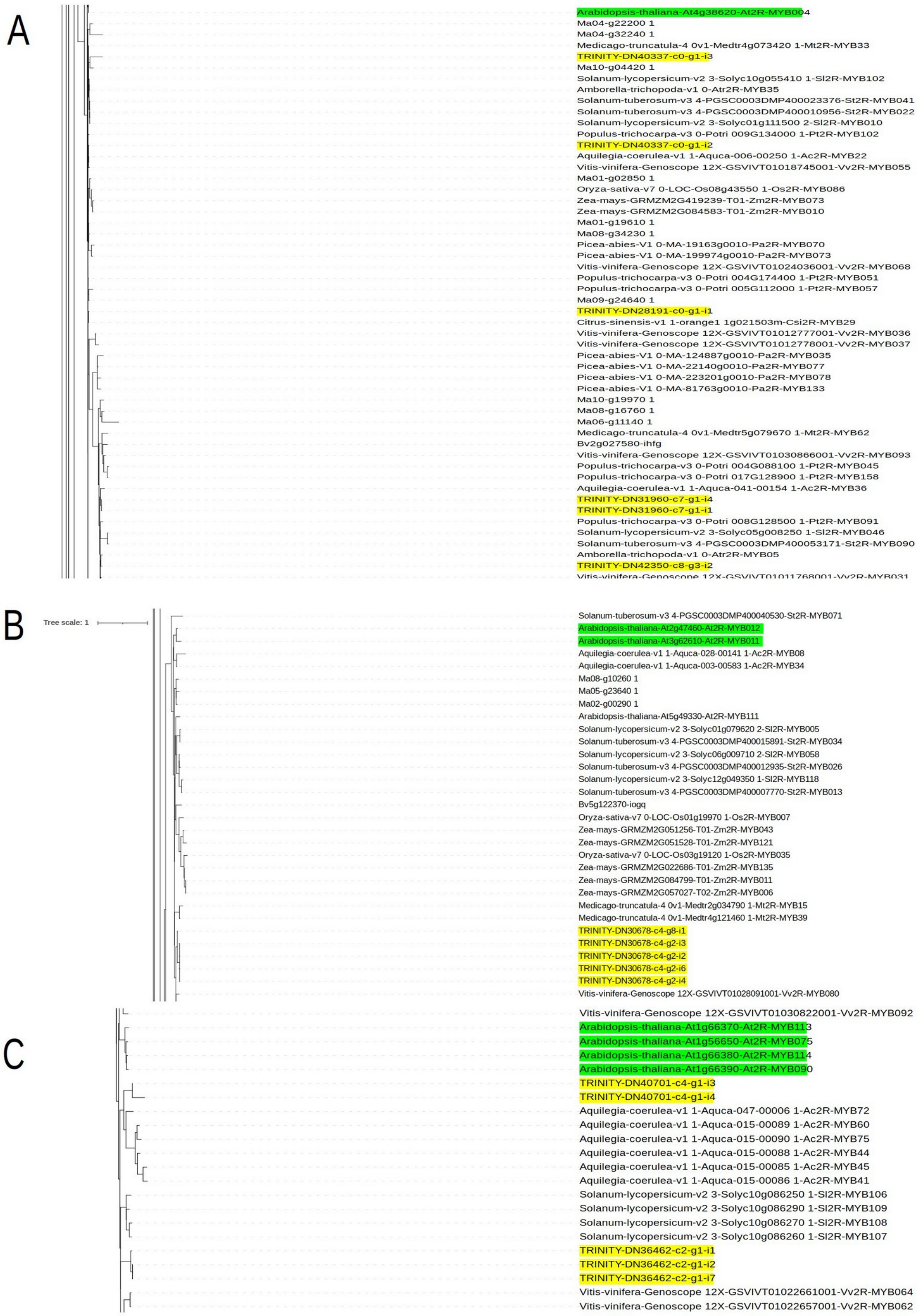


***Figure S8.*** *Phylogenetic trees representing the relationships of MYB transcription factors involved in the flavonoid biosynthesis pathway between Michelia maudiae* (yellow) and corresponding orthologs in *Arabidopsis thaliana* (green). (A) Candidate MYBs related to repressor *MYB4* in *A. thaliana.* (B) Candidate MYBs related to TFs *MYB11 and MYB12* involved in flavonol biosynthesis in *A. thaliana.* (C) Candidate MYBs related to TFs *MYB75*, *MYB90*, *MYB113*, and *MYB114* involved in anthocyanin biosynthesis in *A. thaliana.*

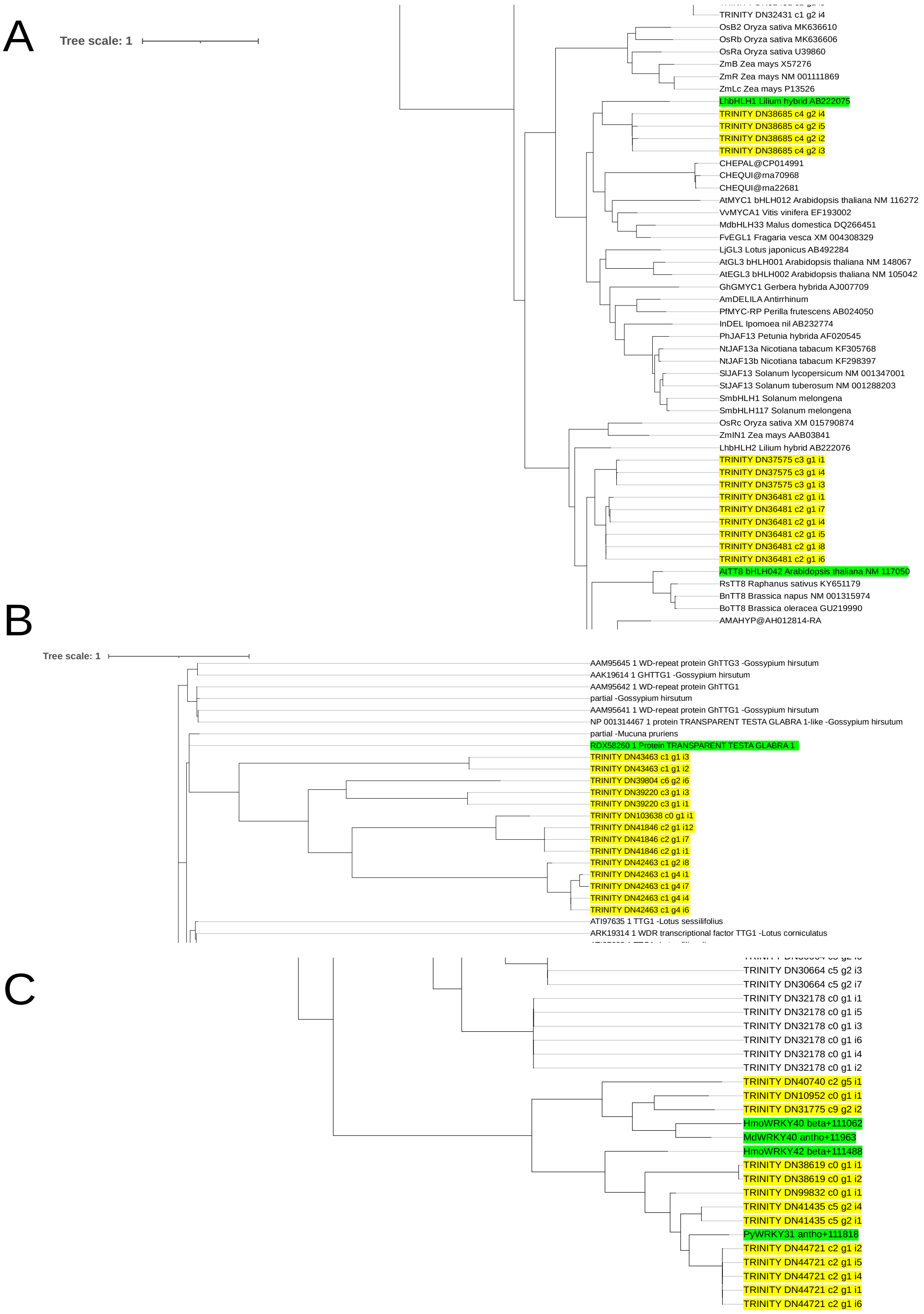


***Figure S9.*** Phylogenetic trees representing the relationships of transcription factors involved in the flavonoid biosynthesis pathway between *Michelia maudiae* (yellow) and corresponding orthologs in other plant species (green). (A) Candidate bHLHs related to transcription factors *bHLH1* in *Lilium spp.*, and *TT8* or *bHLH42* in *A. thaliana* (B) Candidates of TTG1s (C) Candidate WRKYs related to TF WRKY40 involved in anthocyanin biosynthesis in *Malus domestica*, and WRKY31 involved in anthocyanin biosynthesis in *Pyrus spp*.

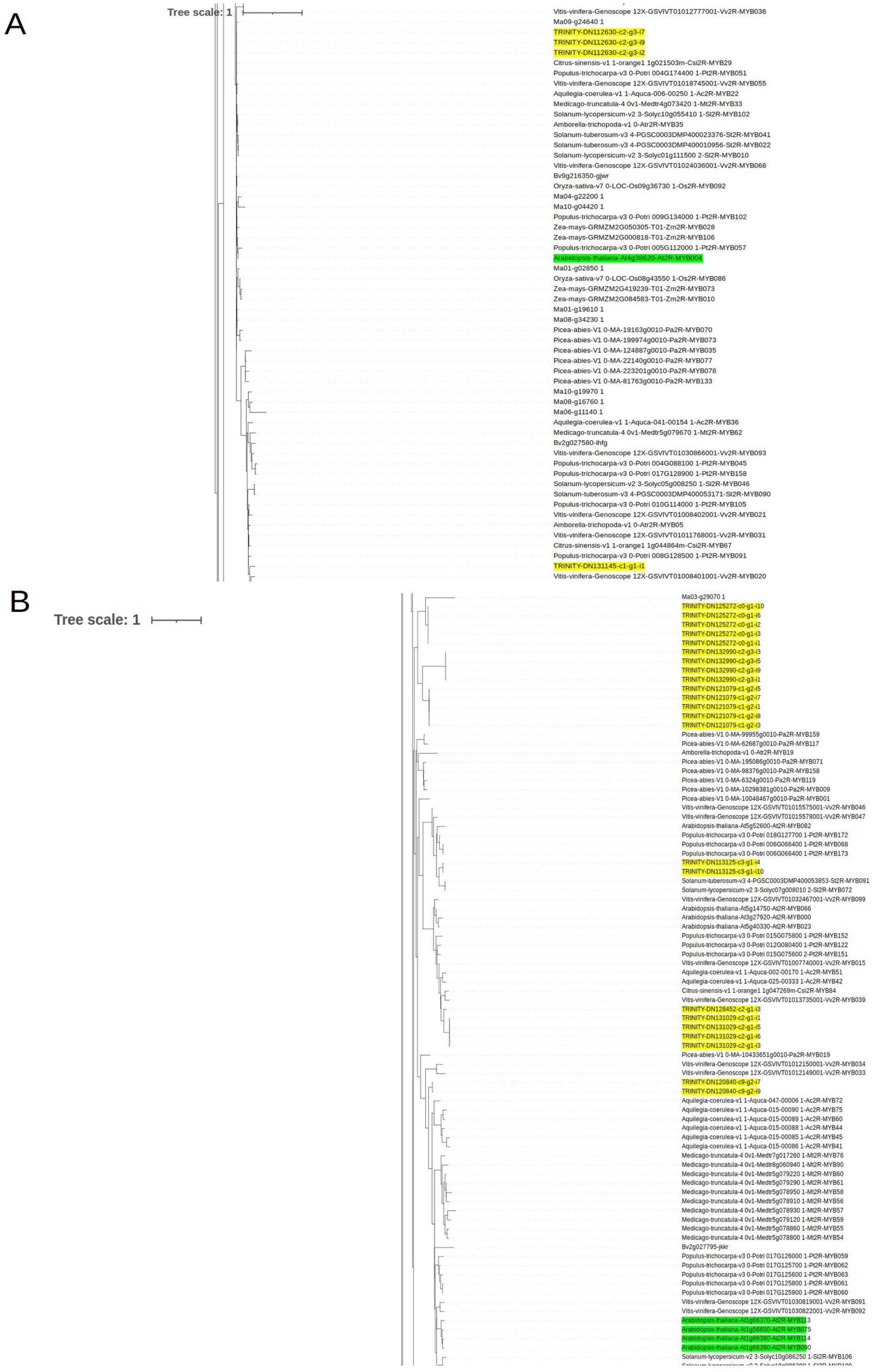


***Figure S10.*** Phylogenetic trees representing the relationships of MYB transcription factors involved in the flavonoid biosynthesis pathway between *Rhododendron obtusum* (yellow) and corresponding orthologs in *Arabidopsis thaliana* (green). (A) Candidate MYBs related to repressor *MYB4* in *A. thaliana*. (B) Candidate MYBs related to TFs *MYB75*, *MYB90*, *MYB113*, and *MYB114* involved in anthocyanin biosynthesis in *A. thaliana*.

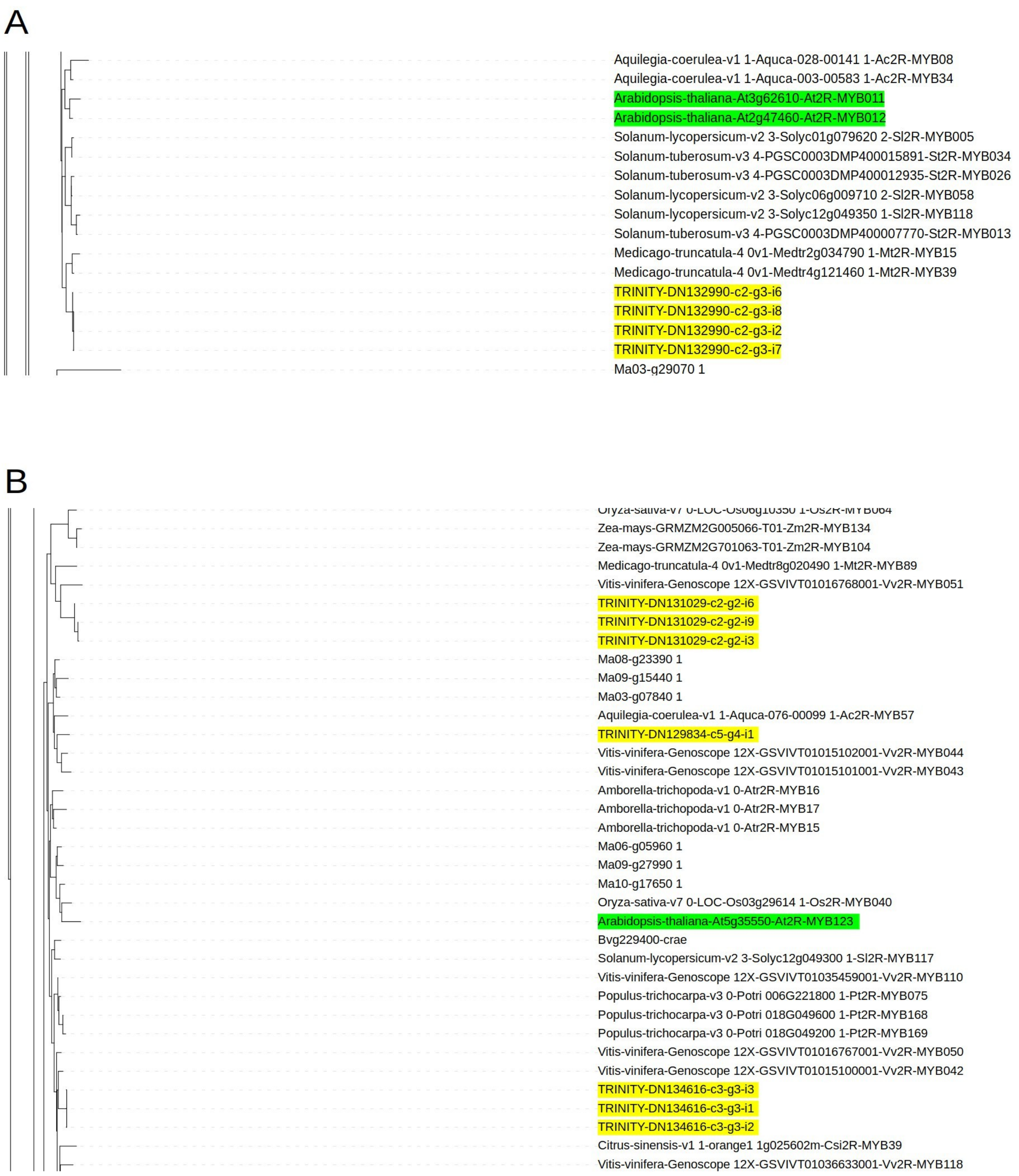


***Figure S11.*** Phylogenetic trees representing the relationships of MYB transcription factors involved in the flavonoid biosynthesis pathway between *Rhododendron obtusum* (yellow) and corresponding orthologs in *Arabidopsis thaliana* (green). (A) Candidate MYBs related to TFs *MYB11 and MYB12* involved in flavonol biosynthesis in *A. thaliana.* (B) Candidate MYBs related to TFs *MYB123* involved in proanthocyanidins biosynthesis in *A. thaliana.*

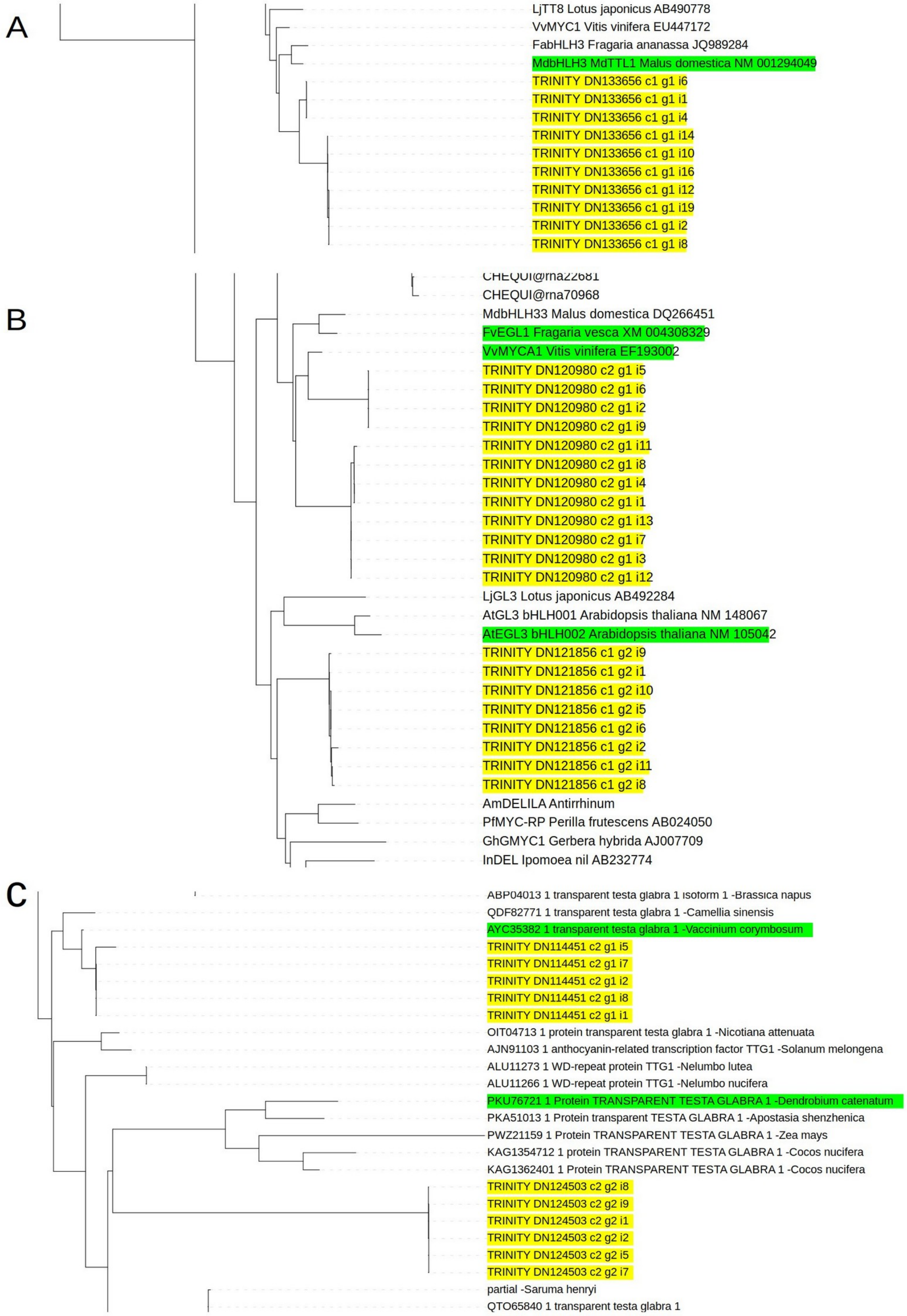


***Figure S12.*** Phylogenetic trees representing the relationships of transcription factors involved in the flavonoid biosynthesis pathway between *Rhododendron obtusum* (yellow) and corresponding orthologs in other plant species (green). (A) Candidate bHLHs related to transcription factor *bHLH3* in *Malus domestica* (B) Candidate bHLHs related to transcription factors *EGL1* in *Fragaria vesca*, *MYCA1* in *Vitis vinifera*, and *EGL3* or *bHLH2* in *A. thaliana* (C) Candidates of TTG1s related to *Vaccinium corymbosum* and *Dendrobium catenatum.*

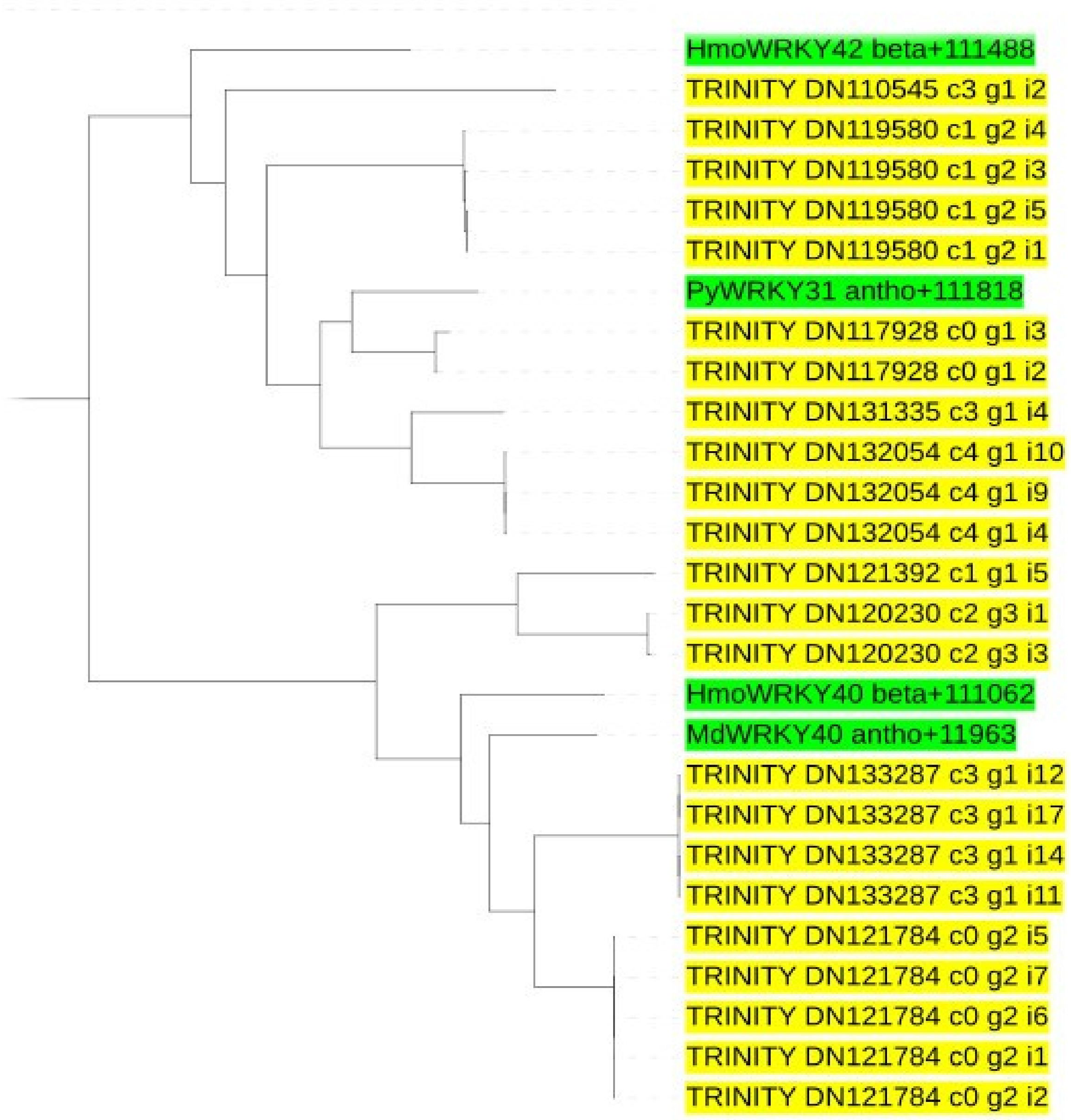


***Figure S13.*** Phylogenetic trees representing the candidate WRKYs in *Rhododendron obtusum* related to TF WRKY40 involved in anthocyanin biosynthesis in *Malus domestica,* and WRKY31 involved in anthocyanin biosynthesis in *Pyrus spp*.

A

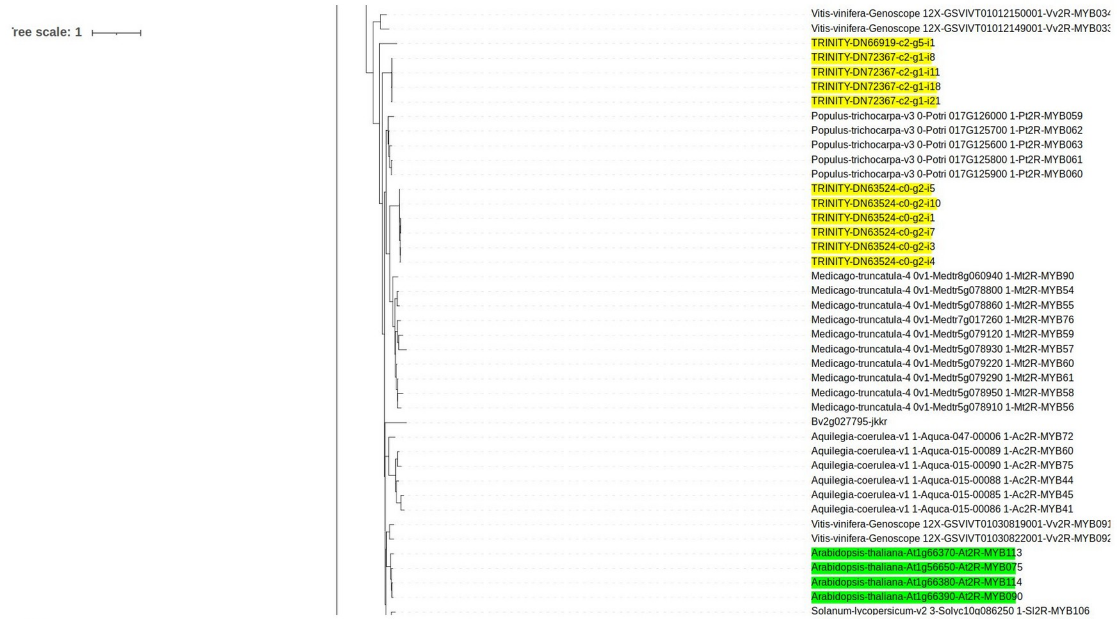


B

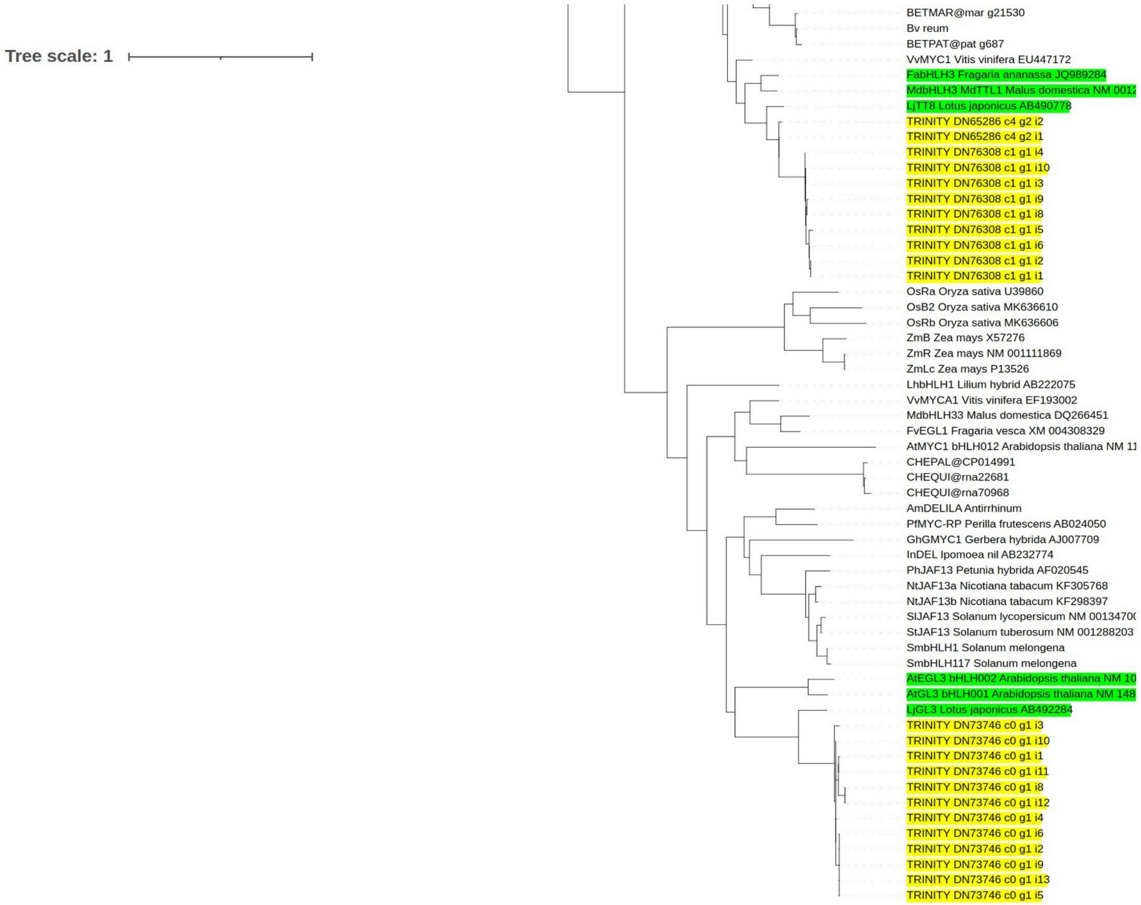


***Figure S14.*** Phylogenetic trees representing the relationships of transcription factors involved in the flavonoid biosynthesis pathway between *Trifolium repens* (yellow) and corresponding orthologs in different species (green). (A) Candidate MYBs related to TFs *MYB75*, *MYB90*, *MYB113*, and *MYB114* involved in anthocyanin biosynthesis in *A. thaliana.* (B) Candidate bHLHs related to transcription factors *bHLH3* in *Malus domestica* and *Fragaria ananassa*, and *TT8* in *Lotus japonicus.*

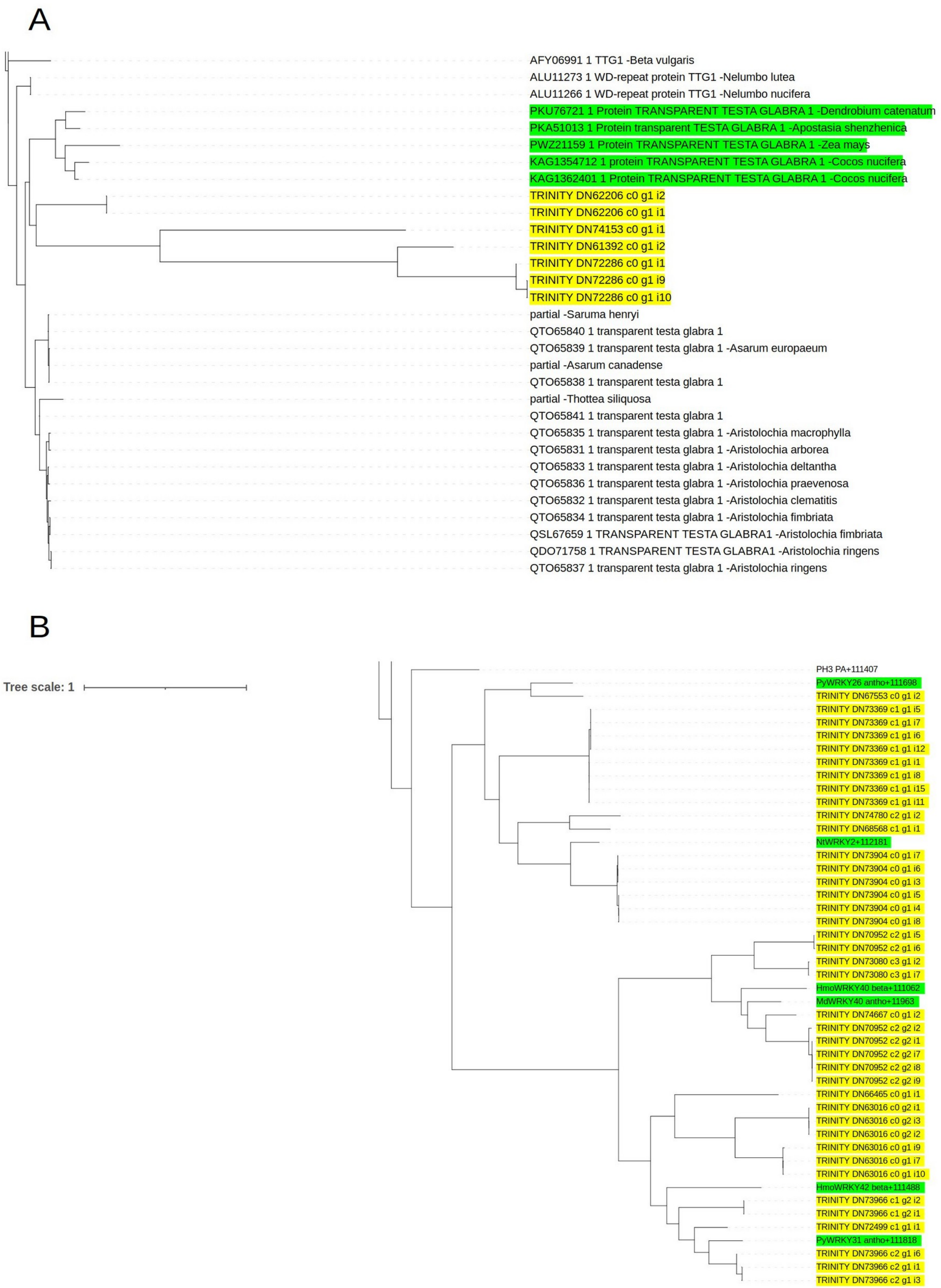


***Figure S15.*** Phylogenetic trees representing the relationships of transcription factors involved in the flavonoid biosynthesis pathway between *Trifolium repens* (yellow) and corresponding orthologs in different species (green). (A) Candidates of TTG1s related to *Dendrobium catenatum, Zea mays* and *Cocos nucifera.* (B) Candidate WRKYs related to TF WRKY40 involved in anthocyanin biosynthesis in *Malus domestica, WRKY2* involved in anthocyanin biosynthesis in *Nicotiana tabacum,* and *WRKY26* and *WRKY31* involved in anthocyanin biosynthesis in *Pyrus spp*.

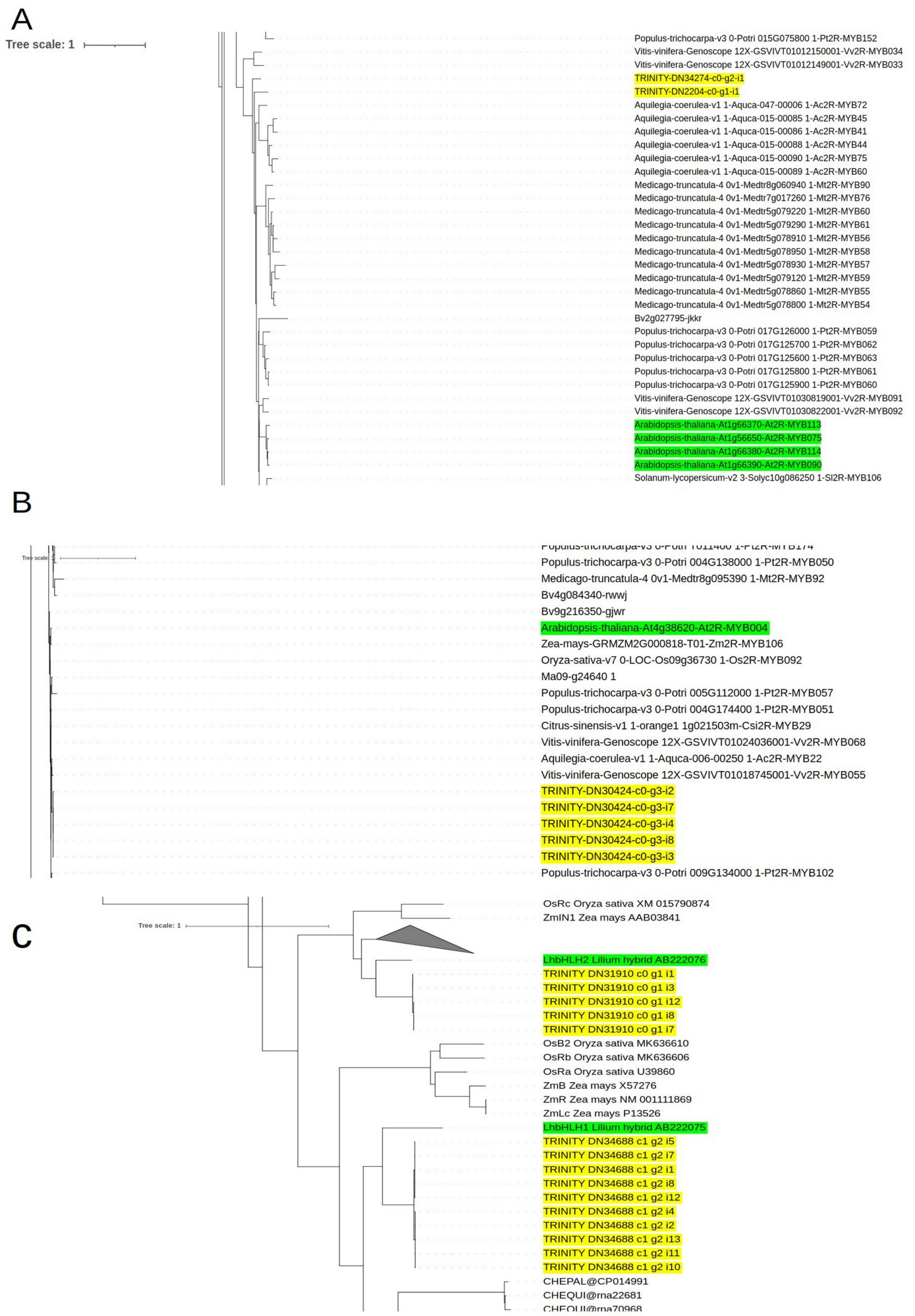


***Figure S16.*** Phylogenetic trees representing the relationships of transcription factors involved in the flavonoid biosynthesis pathway between *Hosta plantaginea* (yellow) and corresponding orthologs in different species (green). (A) Candidate MYBs related to TFs *MYB75*, *MYB90*, *MYB113*, and *MYB114* involved in anthocyanin biosynthesis in *A. thaliana.* (B) Candidate MYBs related to repressor *MYB4* in *A. thaliana.* (C) Candidate bHLHs related to transcription factors *bHLH2 and bHLH1* in *Lilium hybrid.*

A

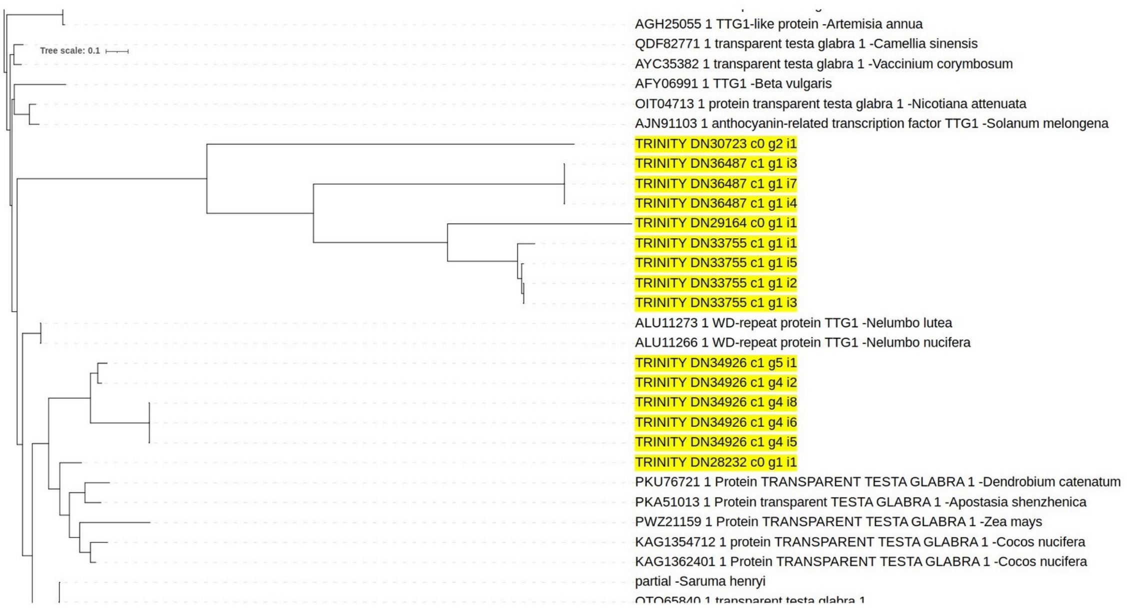


B

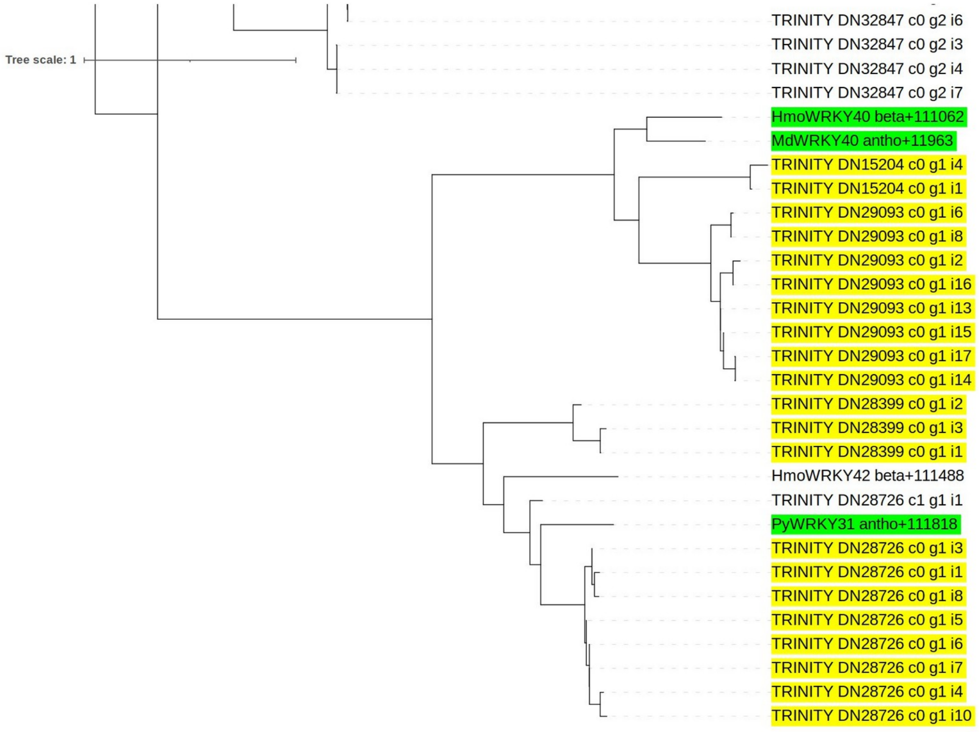


***Figure S17.*** Phylogenetic trees representing the relationships of transcription factors involved in the flavonoid biosynthesis pathway between *Hosta plantaginea* (yellow) and corresponding orthologs in different species (green). (A) Candidates of TTG1s. (B) Candidate WRKYs related to TF WRKY40 involved in anthocyanin biosynthesis in *Malus domestica,* and *WRKY31* involved in anthocyanin biosynthesis in *Pyrus spp*.
